# Systematic comparison of color representations between humans and deep neural networks: towards predicting human color perception in a vast color space

**DOI:** 10.64898/2025.12.10.693611

**Authors:** Nipun Ravindu Wickramanayaka, Masafumi Oizumi

## Abstract

The representational structure of human color perception across a vast color space remains incompletely understood. While classical studies measured numerous color pairs, these measurements compared only similar colors, and exploring the global relationships among thousands of colors has been infeasible due to the time costs of psychophysical experiments. Given these constraints, deep neural networks (DNNs) have attracted attention as a promising tool for providing proxies or predictions of human perception beyond psychophysical experiments. However, it remains unclear which DNNs possess embeddings that geometrically align with human color perception. Furthermore, it is unclear which learning paradigm enables DNNs to acquire a color representation that aligns with that of humans. Here, we systematically investigate which learning paradigm enables DNNs to produce a color representation that is structurally congruent with that of humans, with a focus on three types: self-supervised learning (SSL) that trains on images alone by image reconstruction, supervised learning (SL) that trains on images with category labels, and contrastive language-image pre-training (CLIP) that trains on image-text pairs. We compared the embeddings of DNNs with the human similarity judgments of 93 colors using a rigorous unsupervised method, Gromov-Wasserstein Optimal Transport (GWOT). Our results show that, while each learning paradigm acquires color representations that strongly align with human data at the fine-item level in early layers, only CLIP sustains such a representation at the output. Furthermore, when we leveraged a key advantage of DNNs and investigated the representational structure of 4096 colors, the early layers of each learning paradigm and the output of CLIP consistently converged on their own characteristic structures. These structures present plausible predictions for the large-scale human color representation. Our work demonstrates an approach for exploring unknown territories of human perception using computational models validated in a limited empirical space, and provides predictions for future large-scale psychophysical experiments.

**Author summary:** How do we perceive the vast world of color? Despite extensive research into human color perception, studies evaluating many colors have mostly captured differences between similar colors, while those mapping global relationships are restricted to a few dozen. Consequently, we still do not know the global structure of the massive “color map” that might underlie our perception of thousands of colors, as testing this directly is practically impossible. To explore this space, we turned to deep neural networks, a form of AI. Our first step was to identify models that “see” color in a way that matches humans. We compared models against human data capturing the global relationships among all possible pairs of 93 colors. Using a powerful geometric comparison method, we found the models that matched the human color map. This allowed us to use these models as reliable computational proxies. We then used them to do what human experiments currently cannot: chart a vast global map of 4,096 colors. The human-aligned models consistently converged on two distinct structures. Our work provides the first plausible, testable predictions for the large-scale structure of human color perception and demonstrates a new way to explore otherwise unreachable territories of our perceptual world.

## Introduction

Understanding the detailed representational structure of human color perception is a fundamental challenge in vision and cognitive sciences. While the biological foundations of trichromatic color vision are well-established [1, 2], the precise geometric structure through which humans organize the vast spectrum of perceivable colors remains incompletely understood. Specifically, although previous studies have investigated the geometric structure of human color perception through a variety of psychophysical experiments, including behavioral judgments [3–12], electroencephalography (EEG) [13], magnetoencephalography (MEG) [14], and functional magnetic resonance imaging (fMRI) [15–17], these investigations have typically faced a critical trade-off: studies evaluating a large number of colors have mostly captured local relationships, focusing on perceptual differences between adjacent or similar colors [3, 4, 6, 7], whereas studies aiming to capture global relationships are restricted by the practical limitations of testing human subjects to limited stimuli sets numbering in the dozens [5, 8–17]. A recent study broadened the scale of investigation by successfully mapping the intricate global similarity structure of a 93-color stimulus set that included different tones and achromatic colors [18]; nevertheless, the representational structure for even larger color spaces, such as those spanning thousands of colors, remains largely unexplored.

One powerful methodology for addressing these experimental limitations is the use of deep neural networks (DNNs). DNNs have emerged as powerful tools in computer vision [19–25]; they have achieved remarkable performance across a wide range of perceptual tasks [26–30] and are increasingly regarded as effective computational proxies for the human visual perception [31–36]. In the field of color science, DNNs have also been applied to study diverse phenomena such as color selectivity [37–41], color categorization [42, 43], color constancy [44], color illusions [45], perceptual coherence [46], and the organization of color naming [47]. The use of DNNs enables experiments in ways that are not feasible with human subjects.

However, which DNN models exhibit embeddings whose global geometry aligns with human color perception across a broad color space remains to be established. Previous studies on comparisons of the global geometry of DNN and human color representations have successfully captured correlations using stimulus sets of approximately 20 colors [48–50]. A recent study made progress by comparing a larger set of 93 colors more rigorously, showing that the responses of the state-of-the-art multimodal large language model GPT-4 [51] are consistent with human color similarity judgments [52]. However, the study was unable to clarify whether the internal representations of GPT-4 are naturally consistent with human color perception.

Furthermore, learning paradigms that facilitate the acquisition of a color representation that is geometrically consistent with that of humans have not been identified. Even the study above [52], which provides the most direct insight to date, showed only that the responses of the unimodal large language model GPT-3.5 had limited alignment with human color representation, whereas those of the multimodal large language model GPT-4, which integrates language and vision, achieved remarkable alignment. This finding suggests that language-only models are insufficient, but does not clarify the open question of whether visual information alone is sufficient, or if the integration of visual and linguistic information is necessary.

To address these gaps, our study investigates the contribution of visual information and linguistic grounding to forming a human-like color representation by systematically comparing the embeddings of DNNs trained under distinct learning paradigms. Specifically, we analyze two categories of models: vision-only models and vision-language models. The vision-only models consist of (1) self-supervised learning (SSL) models trained on images alone [53, 54], (2) supervised learning (SL) models trained on images with category labels [25]. The vision-language models are represented by contrastive language–image pretraining (CLIP) models trained on images paired with rich textual descriptions [55], which have been reported to align with human representations across various domains [32, 36, 43]. We compare these models with human similarity judgments of 93 colors [18], representing the largest human-based color similarity dataset among those that captured global relationships across colors by measuring all possible pairs of the colors used in the experiment to date. This comparison allows us to assess which type of learning experience is effective for developing a representational structure that is consistent with that of human color perception, namely the use of visual information alone, visual information with categorical labels, or visual information grounded in rich language.

Of note, a rigorous comparison of these models requires a more sensitive method than standard correlation-based analyses [56–58]. Although such analyses can capture coarse-grained similarities, they are limited by an inherent ambiguity: a high correlation score cannot distinguish whether the alignment is due to a broad, group-to-group correspondence or to a fine-grained, one-to-one correspondence between individual elements [57, 59]. To overcome this limitation and directly assess the fine-grained geometric structure of the representations, we employed Gromov–Wasserstein Optimal Transport (GWOT) [60, 61]. This powerful unsupervised framework allows for sensitive comparison of representational geometry [18, 33, 52, 59, 62–65], making it particularly suitable for detecting subtle differences across models.

Our analysis reveals that, while all three learning paradigms can acquire a color representation that aligns with human similarity judgments at early layers, only the CLIP paradigm sustains this alignment at the output. Notably, this divergence at the output is relatively clearer than that of the supervised comparison, where all three paradigms exhibited substantially high correlations with human data. Furthermore, by exploiting the unique advantage of DNNs to operate beyond human experimental constraints, we probed the geometry of a vast 4096-color representation and found that the second layer embeddings of the three learning paradigms converged upon a relatively simple, undistorted ring-like structure, while the output embeddings of CLIP paradigm converged upon a relatively complex geometry, namely a hollow, distorted ring-like structure with specific protrusions and indentations. These discoveries offer tangible computational predictions for the large-scale organization of human color perception, and pave the way for future, large-scale psychophysical validation. Our “select-and-predict” method thus demonstrates an approach for generating novel and nontrivial hypotheses about human color perception that are presently beyond the reach of direct empirical investigation.

## Methods overview

### Analysis framework

Our study employs a comprehensive analytical framework to systematically compare the structure of color representations between human behavioral data and various deep neural networks (DNNs) in order to probe which learning paradigms enable DNNs to develop internal representations that are structurally analogous to those of human color perception. As briefly illustrated in Figure 1, this process involves two primary stages.

**Fig 1.**
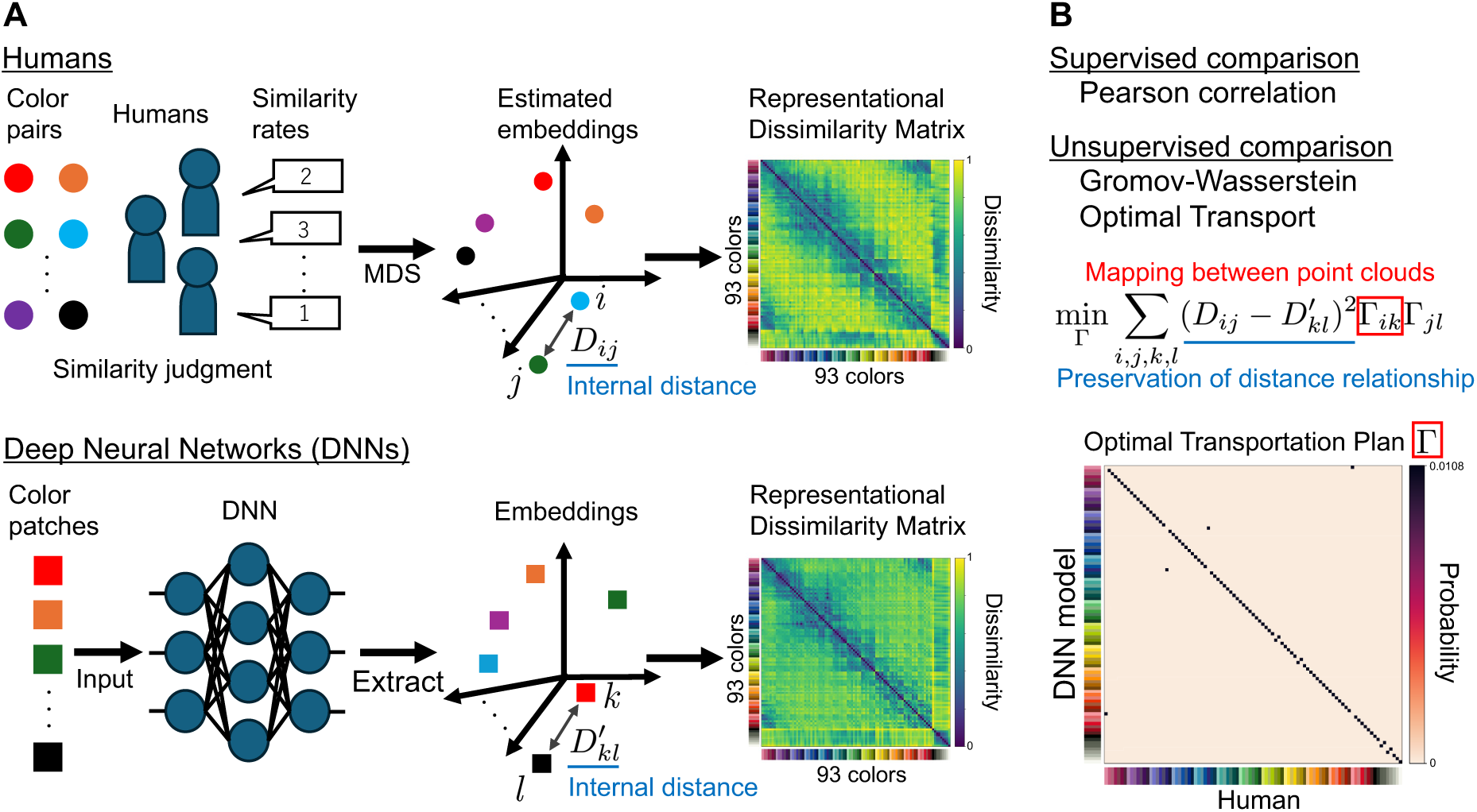
Overview of the analytical framework. (A) Preparation of the representational dissimilarity matrix (RDM). Human data were obtained as large-scale similarity judgments of 93 colors made by human participants in a previous study [18], who rated the similarity of color pairs on a scale from 0 (“very similar”) to 7 (“very different”). Application of multidimensional scaling (MDS) to these ratings produced a set of continuous vector embeddings, which was used to make the human RDM used in this study. For the DNN models, embeddings for the same 93 colors were extracted from the final layer to make the DNN RDM. (B) Comparison of RDM. We compared the RDMs between human and DNNs by two methods, supervised comparison by Pearson correlation and unsupervised comparison by Gromov-Wasserstein Optimal Transport (GWOT).

The first stage is the preparation of representational dissimilarity matrices (RDMs) for both the human behavioral data and the various DNN models, which form the basis of our comparison (Fig. 1A). To establish the human-derived RDM, we first constructed a matrix of the rates from the discrete similarity judgments collected in the previous study [18] for 93 colors selected from the Practical Color Coordinate System (PCCS) [66]. We applied multidimensional scaling (MDS) to this matrix to produce a set of continuous vector embeddings that are directly comparable to the embeddings of DNN, from which we then computed the RDM as the pairwise Euclidean distances between the 93 embeddings. For the DNN-derived RDMs, we first extracted the embeddings for the same 93 colors from models trained under three distinct learning paradigms: self-supervised learning (SSL) trained on images alone; supervised learning (SL) trained on images with category labels; and contrastive language-image pre-training (CLIP) trained on images paired with rich textual descriptions. The RDM for each model was then computed from the pairwise Euclidean distances between these 93 embeddings.

In the second stage, we compared the human-derived RDM with each of the DNN-derived RDMs using two complementary methods (Fig. 1B). The first method is a supervised comparison which is based on a fixed one-to-one correspondence between the colors. For this, we computed the Pearson correlation between the RDMs, known as conventional Representational Similarity Analysis (RSA). We refer to this comparison as supervised because it uses the externally provided identity labels of the colors to fix the one-to-one correspondence in advance. Specifically, each color in the human RDM is matched to the same color in the DNN RDM on the basis of these labels, and the Pearson correlation is then computed between the two RDMs under this label-defined correspondence [59]. This supervised comparison is a standard and widely used approach in neuroscience [56–58], but is limited by an inherent ambiguity: a high correlation score cannot distinguish between a coarse, group-to-group correspondence (in our case, color category to color category alignment) and a fine-grained, one-to-one correspondence (in our case, color to color alignment) [59]. To capture these finer-grained structural properties, our second method is a more rigorous unsupervised comparison using Gromov-Wasserstein Optimal Transport (GWOT) [60, 61]. This method finds an optimal probabilistic mapping, or “transportation plan,” that minimizes the difference between the internal distance structures of two point clouds (in our case, the two sets of color representations ). This alignment process is entirely unsupervised, as it does not use the identity labels of the items being compared (in our case, the colors). This in turn allows for the direct comparison of their intrinsic geometric properties, which cannot be measured by correlation alone. After the alignment is complete, we then quantified the accuracy of the resulting alignment by calculating a “top-1 matching rate” score, which represents the percentage of colors for which the transportation plan correctly assigns the highest mapping probability to its corresponding color in the other structure. This dual-analysis framework provides a multi-faceted evaluation of which learning paradigms produce human-like color representations.

### Human data used in this study

We used human data collected in a previous study [18]. In that study, similarity judgments were collected for all possible pairs of 93 solid-colored circles presented on a uniform grey background. Consequently, the color perception investigated in the present study is explicitly limited to the context of this specific task: judging pure color similarity using uniform patches on a uniform background.

### Selection of DNN models for comparing learning paradigms

To investigate the learning mechanism that enables DNNs to produce internal representations that are consistent with human color perception, this study systematically compares the color representations of DNN models trained under three distinct learning paradigms that represent different types of available information. We evaluated a total of 22 DNN models: two CNN-based and four Transformer-based self-supervised (SSL) models trained on images alone, two CNN-based and three Transformer-based supervised (SL) models trained on images with category labels, and two CNN-based and nine Transformer-based CLIP models trained on images paired with rich textual descriptions. The models are detailed in the Materials and methods section. The results of 93 colors for all 22 models are presented in the Supporting information. An illustrative subset of this larger group are shown in the Results section to facilitate a clear and controlled comparison of the paradigms.

The specific models chosen for this subset were selected to ensure a fair comparison by controlling for the training dataset and varying only the model architecture. For the SSL and SL paradigms, all models featured in the main results were Transformer-based models trained on the ImageNet dataset [67] using the Masked Autoencoder (MAE) objective and a standard supervised object classification, respectively. For the CLIP paradigm, which included models trained on four different datasets (YFCC [68] for CNN-based models and LAION [69], MetaCLIP [70], and DFN [71] for Transformer-based models), we selected the DFN-trained models for the main results. This selection was guided by the results of our unsupervised alignment analysis, in which the DFN-trained models exhibited by far the strongest structural alignment with the human behavioral data. This presentation of the paradigm’s most successful examples facilitates a clear comparison focused on the potential of each learning paradigm, rather than confounding the comparison with less successful instances of the same paradigm. Within each of these selected groups, the main results feature three models that differ only in their architecture size (Vision Transformer Base (ViT-B), Large (ViT-L), and Huge (ViT-H); listed in increasing order of size, to examine whether the alignment with human data differs across architecture sizes).

### Selection of input images for DNN models

For the main analysis, we used uniform color patches as input images to the DNN models. Although such simple inputs may not be ideal for probing the internal representations of complex DNN models in general, they yielded the strongest alignment with human data among the input formats we tested. Specifically, when we used circular color patches on a grey background as input images, the alignment with human data was relatively lower (Fig. S3). Based on these results, we used uniform color patches for the main analysis, since our study aims to identify DNN models whose color representations align with those of humans, and these inputs provided stronger alignment with human data.

### Selection of the dimensionality for MDS visualization

For the visualization of the representational structures using Multidimensional Scaling (MDS), we presented the 2-dimensional results in the main results and the 3-dimensional results in the Supporting information. We chose the 2-dimensional format for the main figures because static 3-dimensional plots can be misleading depending on the viewing angle, making distance relationships difficult to interpret. It should be noted, however, that a 2-dimensional result reflects the compression of one of the axes of the 3-dimensional result. For example, in the human data, the axis related to lightness and chroma was compressed in two dimensions, making two dark colors with opponent hues appear among the most distant from each other (Fig. 2F); in three dimensions, this apparent distance was resolved, and the distance between two dark colors with opponent hues became smaller than that between two light colors with opponent hues (Video S1). To present such richer information, we provided the 3-dimensional MDS results as rotating videos in the Supporting information (Video S1– S6).

**Fig 2.**
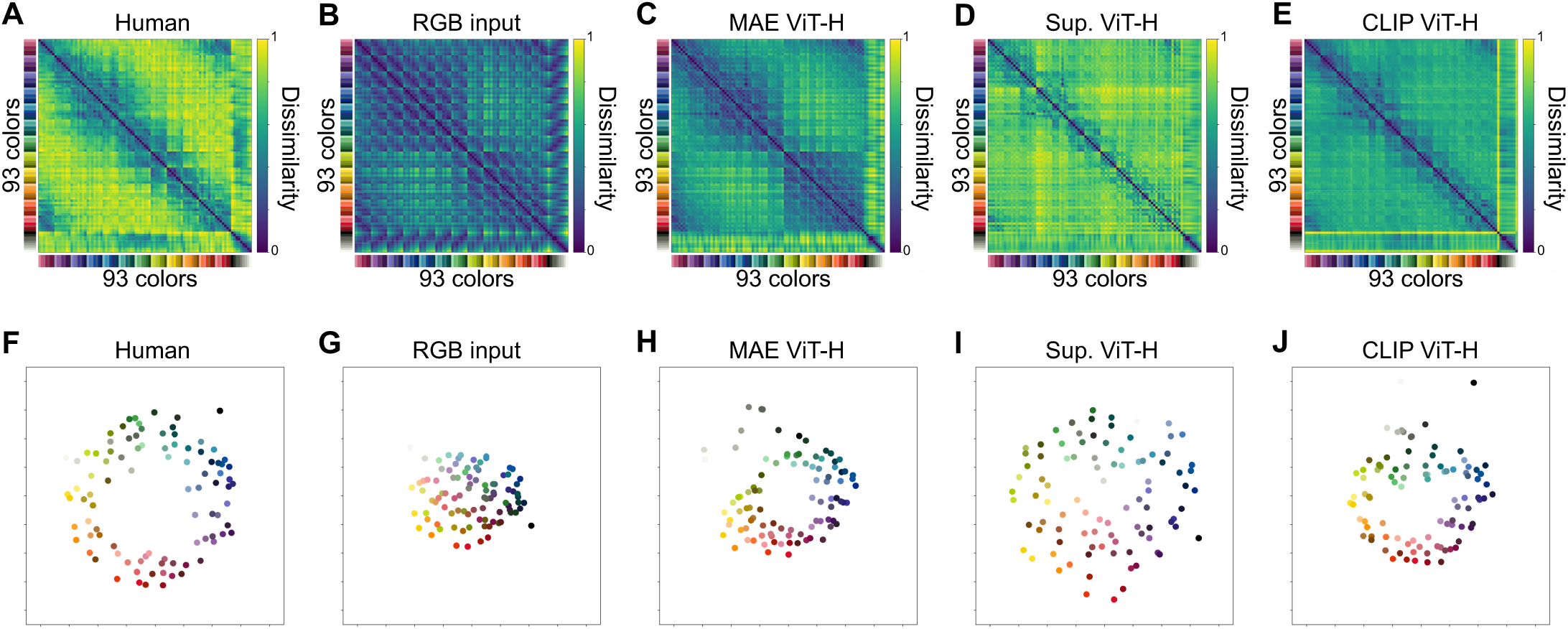
93 colors representation of humans, RGB input, and the output embeddings of the DNN models. (A)-(E) Dissimilarity matrices of 93 colors from (A) the color-neurotypical participants group, (B) the RGB input, (C) a sample from the MAE models, (D) a sample from the SL models, and (E) a sample from the CLIP models. All matrices are normalized to have values between 0 and 1, where 0 means no difference between colors and 1 means the maximum difference for each dissimilarity matrix. (F)-(J) Representation of 93 colors from (F) the color-neurotypical participants group, (G) the RGB input, (H) a sample from the MAE models, (I) a sample from the SL models, and (J) a sample from the CLIP models. All representations are made by applying 2-dimensional MDS on each dissimilarity matrix.

## Results

### Visualization of the color dissimilarity matrices and the color representations of the humans and the output embeddings of DNN models

We first qualitatively compared the structure of color representations through a visual inspection of the representational dissimilarity matrices (RDMs) (Fig. 2A-E). In the human RDM, colors belonging to the same or similar hues are represented as being close to each other, resulting in a block-diagonal-like structure; conversely, colors with dissimilar hues are represented as being far apart (Fig. 2A). The RDMs from all three learning paradigms are visibly distinct not only from the simple structure of the baseline RGB model (Fig. 2B) but also from each other (Fig. 2C-E), indicating that each learning process transforms the input color space to acquire its own unique representational structure. While models from all three paradigms exhibited a block-diagonal-like structure to some extent, visual inspection of the RDMs alone was insufficient to determine which model was most similar to the human data (Fig. 2A-E, S1).

The representational structures showed clearer qualitative similarities and dissimilarities when visualized using 2-dimensional Multidimensional Scaling (MDS), which revealed that the structure of the CLIP paradigm was visually similar to the human data (Fig. 2F-J). The human 93-color representation formed a wide, complex ring structure with intricate tones, and incorporated many achromatic colors (Fig. 2F). In contrast, although the chromatic colors sampled from PCCS form a distinct ring-like for each tone, the RGB baseline resulted in a filled-in structure when all 93 colors, including multiple tones and achromatic colors, were plotted together (Fig. 2G). Among the DNNs, the CLIP model exhibited a wide, complex ring-like structure similar to the human one, with intricate tones and achromatic colors (Fig. 2J). This tendency was consistent regardless of architectural size (Fig. S2I-K). The MAE model also formed a wide, complex ring-like structure with intricate chromatic colors (Fig. 2H), but this trend was dependent on architectural size (Fig. S2C-E). In contrast to these paradigms, the SL models failed to form a clear ring-like structure (Fig. 2I, S2F-H). To better understand these qualitative findings of varying degrees of similarity across the human and model representations, we conducted a quantitative analysis to objectively assess the degree of similarity between the representational structures.

### Correlations between the color representations of humans and DNN models

A quantitative analysis using Pearson correlation revealed that most DNN models exhibited higher correlations with the human RDM than the RGB baseline at early layers, while their behavior at subsequent layers differed depending on the learning paradigm (Fig. 3). Notably, even the baseline RGB model, which represents the input colors prior to any learning, showed a substantial correlation with the human RDM (*ρ* = 0.64). All featured DNN models surpassed this baseline score at early layers. At subsequent layers, the three learning paradigms exhibited the following overall patterns, with some variation across architecture sizes: MAE models exhibited a drop in correlation at an early stage, followed by a recovery near the output; SL models exhibited a generally continuous decline in correlation; and CLIP models exhibited a drop in correlation at relatively late layers, followed by a recovery near the output. When circular color patches were used as input images instead of uniform color patches, however, the correlation scores were essentially inverted in the intermediate layers: the CLIP model was almost consistently above the other two models with uniform color patches, whereas the SL model was almost consistently above the other two models with circular color patches, with which the correlation scores of the MAE and CLIP models were on average lower (Fig. S3E-H).

**Fig 3.**
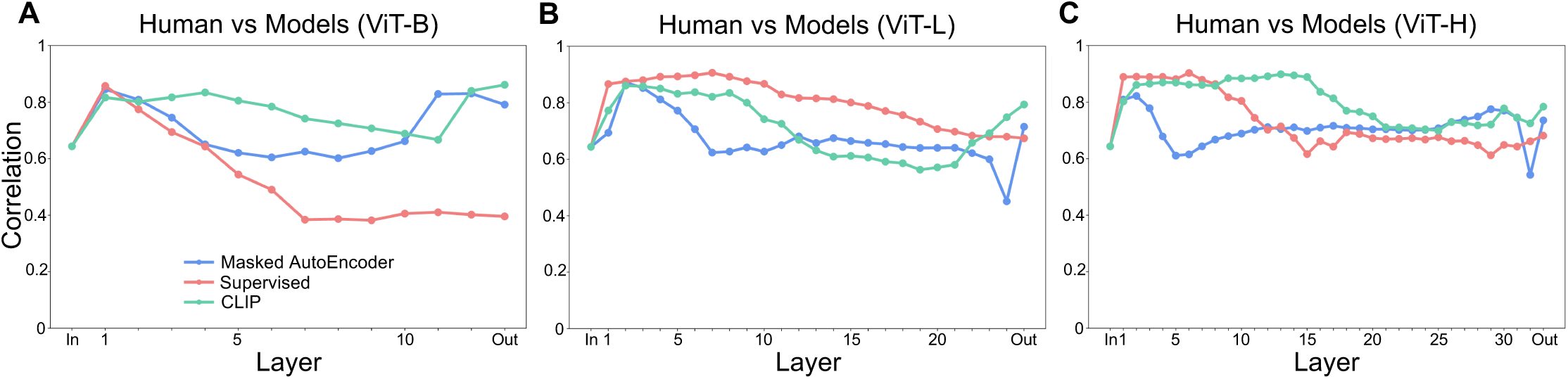
Correlations of color similarity structures for 93 colors between humans and the DNN models. Quantitative comparison based on Pearson correlations between the dissimilarity matrices of humans and the embeddings of each layer of the DNN models for (A) ViT-B, (B) ViT-L, and (C) ViT-H architectures. In all panels, “In” and “Out” on the horizontal axis denote the RGB input and the output embedding, respectively.

### A CLIP model aligns with humans in an unsupervised manner

While the preceding correlation-based analysis revealed high correlation scores for all DNN models, this finding is limited by an inherent ambiguity; it cannot distinguish whether the alignment is due to a coarse, group-to-group correspondence (in our case, alignment of colors within the same hue as represented in the block diagonal structure in RDMs) or a fine-grained, one-to-one correspondence (in our case, same color-to-same color alignment) [57, 59]. To resolve this ambiguity, we directly assess their geometric structure using the unsupervised alignment method, Gromov-Wasserstein Optimal Transport (GWOT) [60, 61]. Before presenting a systematic comparison across all models, we first illustrate the principles of this more sensitive analysis using a single, representative example: alignment of the human RDM (Fig. 4A) with that from a high-performing CLIP model (Fig. 4B).

**Fig 4.**
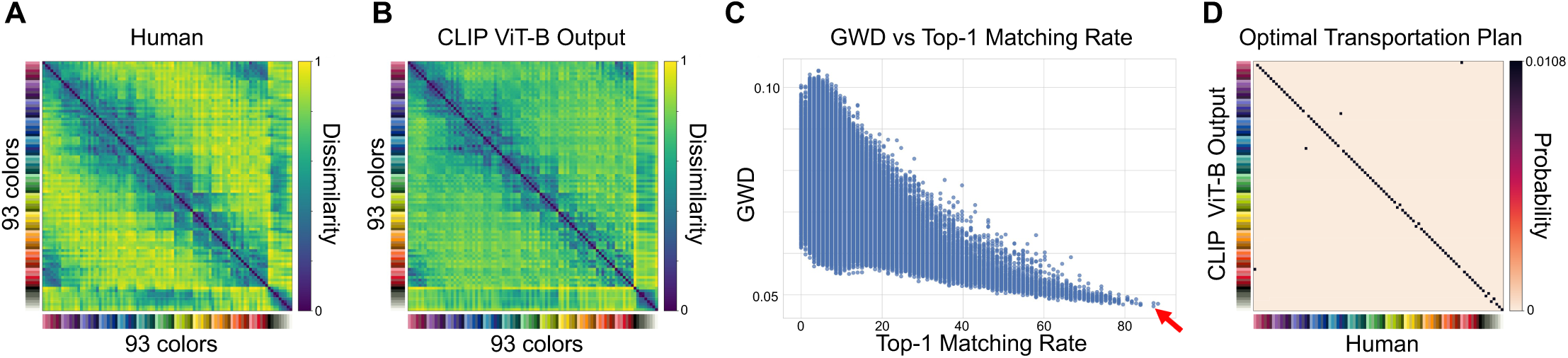
Unsupervised alignment of color similarity structures for 93 colors between humans and the output embeddings of a CLIP model based on GWOT. (A)-(B) Dissimilarity matrices of 93 colors from (A) the color-neurotypical participants group, and (B) a sample from the CLIP model. All matrices are normalized to have values between 0 and 1, where 0 means no difference between colors and 1 means the maximum difference for each dissimilarity matrix. (C) Optimization results over 100,000 iterations with different initial values. The red arrow indicates the result with the lowest GW distance, whose transportation plan is shown in (D). (D) Optimal transportation plan between the dissimilarity matrices of human behavior and a CLIP model.

The goal of GWOT is to find an optimal mapping, or the “optimal transportation plan”, that minimizes the structural difference between two point clouds, quantified as the Gromov-Wasserstein Distance (GWD). Two RDMs (Fig. 4A and B) are used as inputs for finding the optimal transportation plan. As this optimization is non-convex, we ran the algorithm multiple times from different random initializations and selected the solution that achieved the lowest GWD (Fig. 4C), which corresponds to the optimal transportation plan shown in Fig. 4D.

The GWOT analysis of this representative CLIP model revealed a high degree of structural congruence with the human data. The resulting optimal transportation (OT) plan, shown in Fig. 4D, visualizes the probabilistic mapping between the 93 colors of the human similarity structure and those of the CLIP model, exhibiting a remarkable concentration of probability mass along its main diagonal. This diagonal concentration indicates that for most colors, the highest mapping probability is assigned between the corresponding colors in the human and the CLIP model similarity structures. This correct one-to-one correspondence (top-1 matching rate) was achieved for 87.1% of the colors, even though the alignment was performed without any supervised information about color identities. Achievement of such a high matching rate via an unsupervised method strongly suggests that the intrinsic geometric structure of the CLIP model’s color representation is highly consistent with that of human color perception. This example therefore demonstrates that GWOT can assess a detailed structural similarity that cannot be captured by correlation-based measures alone. We now apply this analysis to systematically compare all models.

### Unsupervised alignment between humans and DNN models

Having illustrated the principle of the GWOT analysis with a representative example, we now present the results of applying this method systematically to compare the human similarity structure with that of all DNN models and the RGB baseline (Fig. 5).

**Fig 5.**
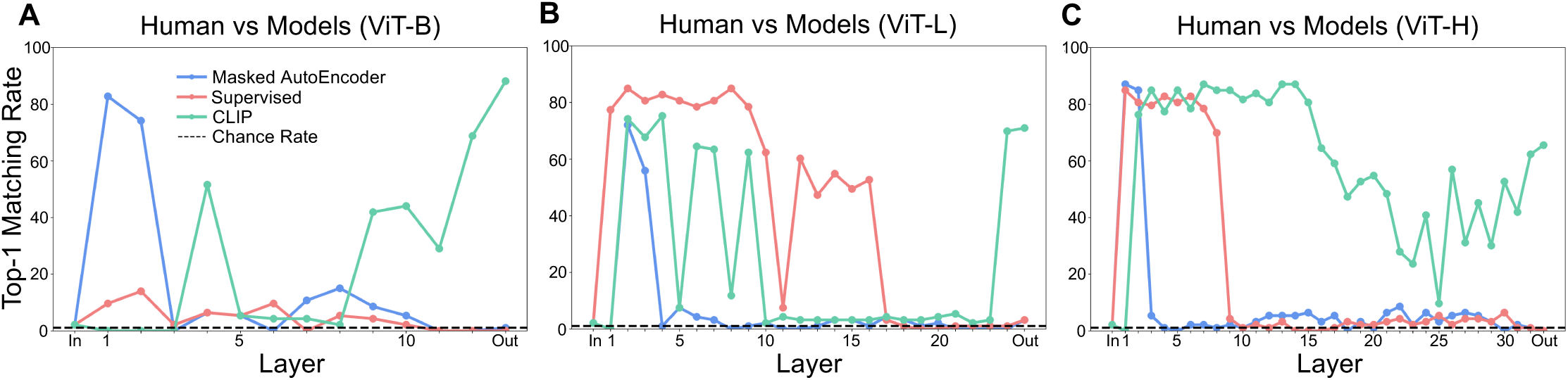
Unsupervised alignment of color similarity structures for 93 colors between humans and the DNN models based on GWOT. Top-1 matching rates of unsupervised alignment based on GWOT between the dissimilarity matrices of humans and the embeddings of each layer of the DNN models for (A) ViT-B, (B) ViT-L, and (C) ViT-H architectures. In all panels, “In” and “Out” on the horizontal axis denote the RGB input and the output embedding, respectively.Analysis of consistency across the embeddings of second layer and output of DNN models

First, the RGB input achieved a top-1 matching rate of only 2.15% (Fig. 5) (or 3.23% even with a tenfold increase in the number of GWOT initializations (Fig. S4B)), which is barely above the theoretical chance rate (1/93 *≈* 1.08%, shown as a dashed line in the figure). This result clearly exemplifies that a high correlation score does not guarantee fine-grained geometric alignment, as this model’s extremely low matching rate was observed alongside its substantial correlation (*ρ* = 0.64) in the supervised analysis (Fig. 3).

In striking contrast, all three learning paradigms included architectures that achieved high top-1 matching rates at early or intermediate layers. Specifically, MAE models achieved scores exceeding 70% at early layers across all architecture sizes (Fig. 5). SL models achieved scores around 80% at early layers in ViT-L and ViT-H (Fig. 5B, C), and scores around 60% at intermediate layers in ViT-L (Fig. 5B). CLIP models achieved scores exceeding 60% at early layers in ViT-L, and scores around 80% from early to intermediate layers in ViT-H (Fig. 5B, C). A similar pattern was also observed in CNN-based models, where all three learning paradigms achieved high top-1 matching rates at intermediate layers (Fig. S5C, D). Of note, these top-1 matching rates achieved by the DNN models are higher than those achieved by the advanced color space models (Fig. S6).

At the output, however, only the CLIP paradigm sustained high structural alignment with human data. Specifically, the output embeddings of all featured CLIP models achieved top-1 matching rates exceeding 60%, with the top-performing ViT-B model reaching 88.2% (Fig. 5) (or 87.1% with a tenfold increase in the number of GWOT initializations (Fig. S4B)). Although this alignment depended on the training dataset and architecture size (Fig. S4B), the output embeddings of the CLIP paradigm nevertheless achieved markedly higher matching rates than those of the other paradigms. Specifically, the matching accuracies of the output embeddings of MAE and SL paradigms did not substantially differ from the chance rate (Fig. 5). A similar pattern was observed in CNN-based models, except that one self-supervised learning model exhibited some degree of alignment at the output (Fig. S5C), although the matching rate was lower than those at its own intermediate layers (Fig. S5C) and also lower than that of the CLIP model that aligned with human data at the output (Fig. S5D). This loss of alignment at the output is particularly noteworthy given that these embeddings exhibited high correlation scores in the supervised comparison (Fig. 3, S5A, B). Of note, the top-1 matching rates were on average lower when circular color patches were used as input images than when uniform color patches were used, particularly those of the CLIP model and particularly at the output (Fig. S3I-L).

### Complementary roles of the supervised and unsupervised comparisons

A side-by-side examination of the Pearson correlation (Fig. 3) and the GWOT-based top-1 matching rates (Fig. 5) indicates that the supervised and unsupervised comparisons provide complementary information about the alignment between the color representations of humans and DNNs. On the one hand, the supervised comparison based on correlation reveals gradual differences that the unsupervised comparison did not discern. Since the top-1 matching rate reflects whether the fine-grained, one-to-one correspondence is achieved, a low matching rate does not necessarily mean that a representation is far from the human color similarity structure. In addition, because the color similarity structures of both humans and the DNN models are approximately symmetric with respect to rotation and reflection, GWOT can converge to flipped (Fig. S7H) or rotated solutions (Fig. S7E, M, T) at some layers, so that the top-1 matching rate can change sharply even between layers across which the correlation changes only gradually. The top-1 matching rate can also decrease at layers where GWOT converges to locally shuffled solutions (Fig. S7Q, R, V, X). For layers where the top-1 matching rate remains near the chance rate, or for layers where it fluctuates sharply (Fig. 5, S5C, D), the correlation scores reveal, in a more gradual manner, how closely each representation resembles the human color similarity structure relative to the RGB baseline (Fig. 3, S5A, B). On the other hand, the unsupervised comparison based on GWOT reveals fine-grained, one-to-one structural alignment that the supervised comparison cannot discern. While all three learning paradigms included architectures that aligned with human data at early or intermediate layers, only the CLIP paradigm sustained this alignment at the output (Fig. 5, S4B, S5C, D). This divergence was not apparent in the supervised comparison, where the output embeddings of all three paradigms exhibited substantially high correlations with human data (Fig. 3, S4A, S5A, B). Rather than one being superior to the other, the two comparisons are complementary, and combining them provides a more complete characterization of the alignment between the color representations of humans and DNNs than either could alone.

### Analysis of consistency across the embeddings of second layer and output of DNN models

This section presents a direct comparison between the embeddings of each learning paradigm that geometrically aligned with human data to assess whether these human-aligned embeddings share the same geometric structure with each other. This analysis is driven by two primary motivations arising from our preceding human-vs-model results (Fig. 5). The first motivation is to investigate the consistency among the early-layer embeddings of the three learning paradigms. Our finding that the early-layer embeddings of all three learning paradigms aligned with human data (Fig. 5) led us to ask whether these embeddings also align with each other. The second motivation is to assess the consistency between the output embeddings of CLIP and the early-layer embeddings of each learning paradigm. When measured by alignment with human data, the output embeddings of CLIP and the early-layer embeddings of each learning paradigm, including CLIP itself, showed comparable scores in the preceding analysis (Fig. 5). Building on this observation, we sought to test whether they are structurally aligned with each other. To address these motivations, we used the second-layer embeddings as a representative of the early-layer embeddings and compared them with the output embeddings of each learning paradigm.

Our analysis of consistency among the second-layer embeddings of each learning paradigm revealed that they exhibited high structural alignment with each other (Fig. 6B). Specifically, the second-layer embeddings that aligned with human data showed very high top-1 matching rates with each other, with some pairs reaching 100% (Fig. 6B). These pairs also exhibited correspondingly high correlations with each other (Fig. 6A). These results indicate that Transformer-based DNNs acquire a common color representation in their early layers within the range of the 93 PCCS colors regardless of the learning paradigm, which is also qualitatively shown by the visualization of the representations (Fig. S8, S9).

**Fig 6.**
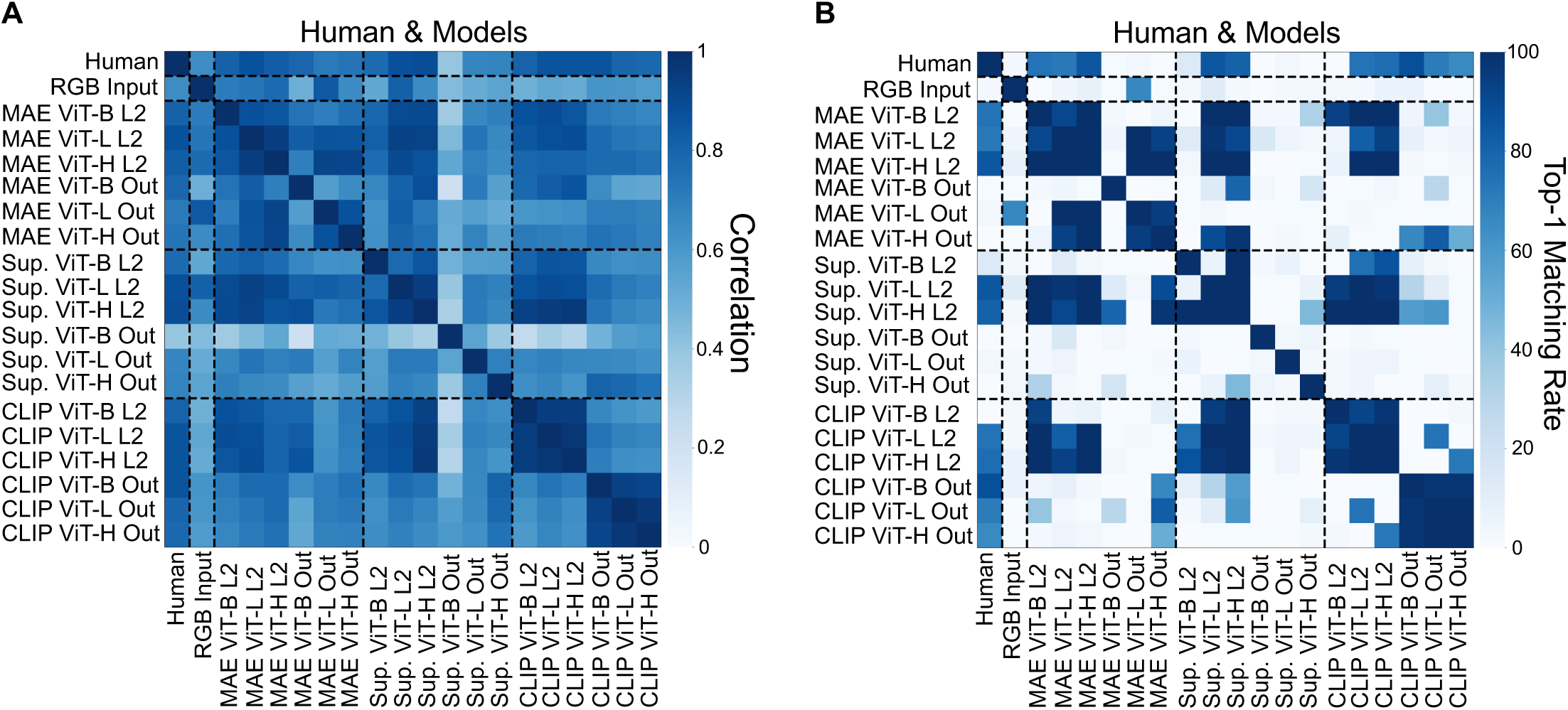
Supervised and unsupervised comparison of color similarity structures for 93 colors between each pair of humans, RGB input, and the embeddings of the second layer and the output of the DNN models. (A) Supervised comparison based on Pearson correlations between each pair of humans, RGB input, and the embeddings of the second layer and the output of DNN models. (B) Unsupervised comparison based on GWOT between each pair of humans, RGB input, and the embeddings of the second layer and the output of DNN models. In each panel, “L2” and “Out” in the labels denote the embeddings of the second layer and the embeddings of the output, respectively.

Our analysis of consistency between the output embeddings of CLIP and the second-layer embeddings of each learning paradigm revealed that the output embeddings of CLIP exhibited little structural alignment with the second-layer embeddings of any learning paradigm, including CLIP itself (Fig. 6B). Specifically, the output embeddings of CLIP showed some degree of top-1 matching rate with the second-layer embeddings of CLIP at the same architecture size and with those of some other learning paradigms, but overall the top-1 matching rates were very low (Fig. 6B). The correlations between the output embeddings of CLIP and the second-layer embeddings of each learning paradigm, including CLIP itself, were also lower than the correlations among the second-layer embeddings (Fig. 6A). Together with the finding that the output embeddings of CLIP exhibited high structural alignment with each other (Fig. 6, S10, S11), these results indicate that the output embeddings of CLIP acquire a color representation with a unique geometric structure distinct from that of the early-layer embeddings of any learning paradigm, including CLIP itself.

### Beyond human experimentation: probing the vast 4096-color representations in DNNs

Having identified the embeddings of DNN models that geometrically align with human color perception using the 93 PCCS colors, we now leverage a key advantage of computational models to explore a vastly larger color space that is inaccessible to human experimentation. For this analysis, we created a 4096-color set by uniformly sampling 16 values from each of the R, G, and B channels (16^3^ = 4096), primarily because RGB serves as the direct input space for the evaluated DNN models. This sampling method provides a more uniform and dense coverage of the entire digital RGB cube, in contrast to the PCCS’s perceptually-structured sampling based on hue and tone. This allows us not only to test the robustness and generalizability of the learned representations by DNNs but also to use the high-performing models to make novel predictions about the large-scale structure of human color perception.

### Visualization of the 4096-color representations of the embeddings of the second layer and the output of DNN models

A qualitative analysis of the 4096-color representations, visualized using 2-dimensional MDS, reveals that the second-layer embeddings of all three learning paradigms exhibited a relatively undistorted ring-like structure, while the output embeddings differed across the learning paradigms, with the CLIP models appearing to converge on a relatively complex geometry (Fig. 7). As a baseline reference, the RGB model forms a filled-in, cube-like structure (Fig. 7A). This is a trivial outcome given that the colors were uniformly sampled from the raw three-dimensional input space. In clear distinction to this baseline, the second-layer embeddings of all three learning paradigms generally exhibited a relatively undistorted ring-like structure, although there was some variation in the specific shape across paradigms and some architectures yielded more filled-in point clouds (Fig. 7B-D, H-J, N-P). In contrast, the output embeddings differed across the learning paradigms. While the output embeddings of MAE and SL models exhibited different shapes across architecture sizes (Fig. 7E-G, K-M), the CLIP models consistently produced a comparable geometric structure, namely a wide, hollow, distorted ring-like shape with specific protrusions and indentations to which the achromatic colors were generally located externally (Fig. 7Q-S). These qualitative results for the output embeddings were also observed when the representations were visualized in three dimensions (Video S6).

**Fig 7.**
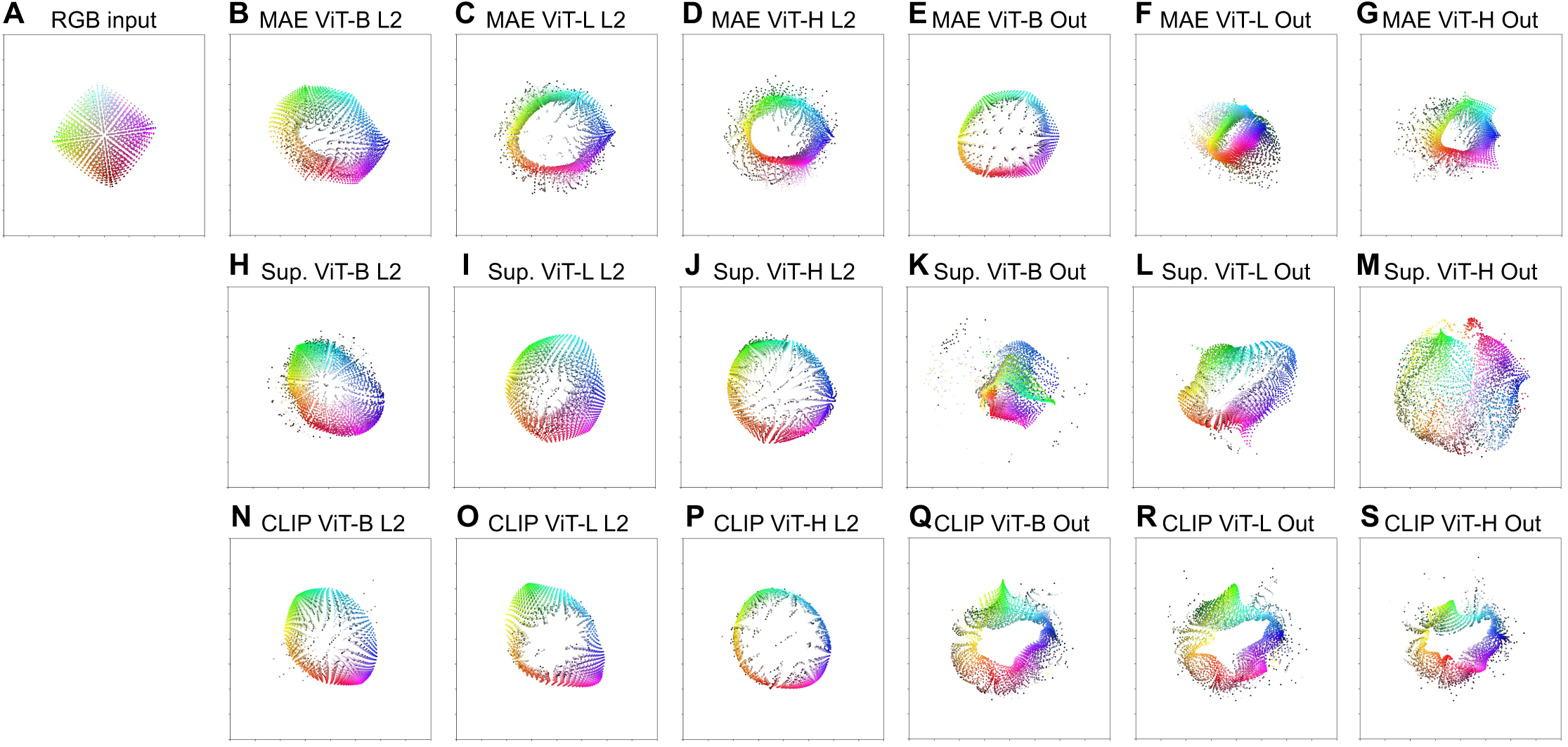
4096 colors representation of RGB input and the embeddings of the second layer and the output of the DNN models. Representation of the 4096 colors from (A) the RGB input, (B)-(D) the embeddings of the second layer of the MAE models, (E)-(G) the embeddings of the output of the MAE models, (H)-(J) the embeddings of the second layer of the SL models, (K)-(M) the embeddings of the output of the SL models, (N)-(P) the embeddings of the second layer of the CLIP models, (Q)-(S) the embeddings of the output of the CLIP models. In all panels, “L2” and “Out” in the labels denote the embeddings of the second layer and the embeddings of the output, respectively. All representations are made by applying 2-dimensional MDS on the respective dissimilarity matrix.

### Correlations and top-1 matching rates between 4096-color representations of the embeddings of the second layer and the output of DNN models

Quantitative analysis of the measured correlations and top-1 matching rates between 4096-color representations provided objective evidence for the high consistency of the output embeddings of the CLIP models and the second-layer embeddings of the three learning paradigms (Fig. 8). Specifically, the output embeddings of the CLIP model with the smallest architecture (ViT-B) exhibited moderate top-1 matching rates with the output embeddings of the CLIP models with the larger architectures (ViT-L and ViT-H) (Fig. 8B). Similarly, the second-layer embeddings of the MAE model with the largest architecture (ViT-H) exhibited moderate top-1 matching rates with the second-layer embeddings of the MAE models with the smaller architectures (ViT-L and ViT-B) (Fig. 8B). In addition, certain levels of top-1 matching rates was observed among the second-layer embeddings of CLIP models, between the output embeddings of the MAE models with the larger architectures (ViT-L and ViT-H), and between the second-layer embeddings of different learning paradigms (Fig. 8B). Although numerically smaller than the highest rates in the 93-color analysis, these results are arguably more significant given the combinatorial magnitude of the search space, comprising 4096! (*>* 10^10,000^) possible permutations. As for their correlations, pairs with high top-1 matching rates also showed high correlations, consistent with the results for the 93-color analysis (Fig. 8A). In addition, correlations among the second-layer embeddings were relatively higher than those observed in the 93-color analysis, further supporting their consistency across models and learning paradigms.

**Fig 8.**
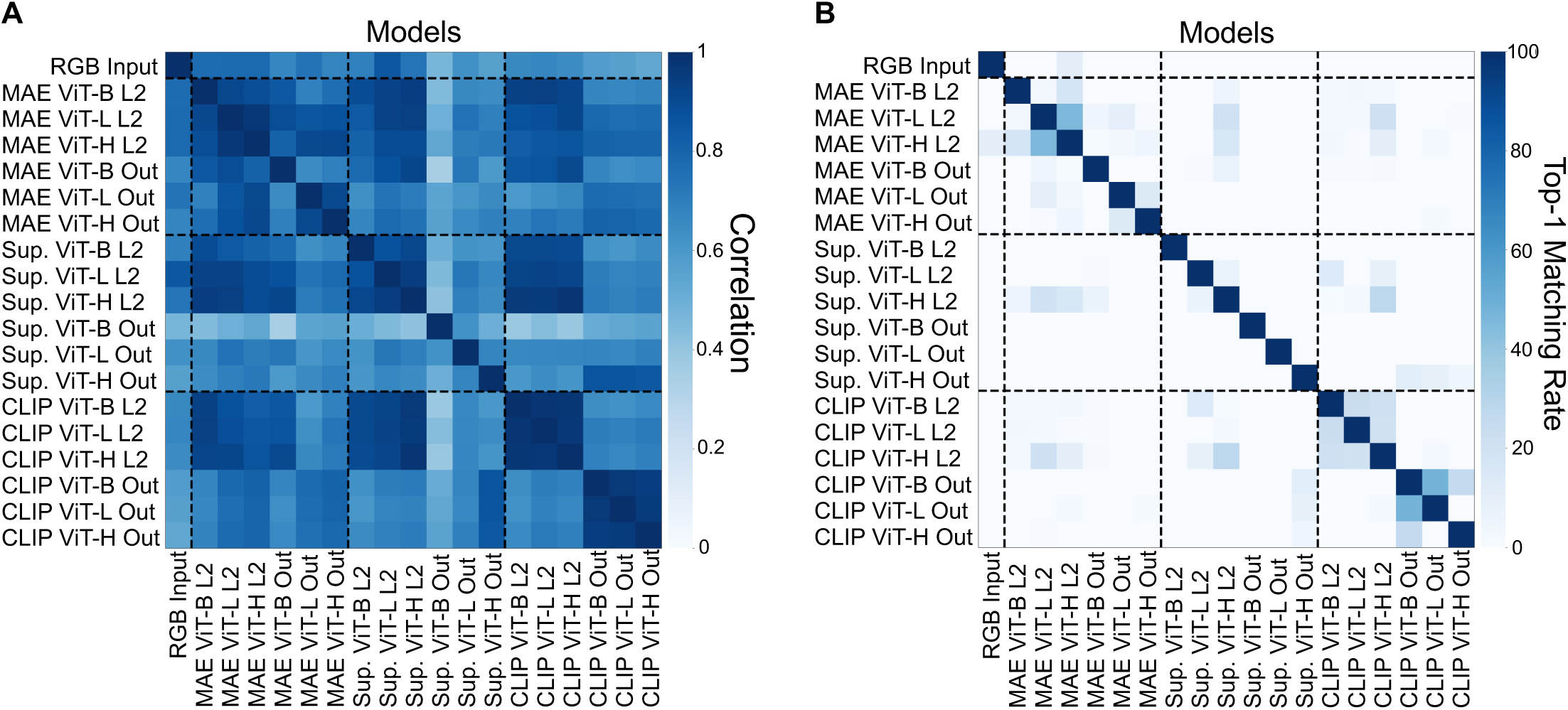
Supervised and unsupervised comparison of color similarity structures for 4096 colors between each pair of RGB input and the embeddings of the second layer and the output of the DNN models. (A) Supervised comparison based of Pearson correlation between each pair of RGB input and the embeddings of the second layer and the output of DNN models. (B) Unsupervised comparison based on GWOT between each pair of RGB input and the embeddings of the second layer and the output of DNN models. In all panels, “L2” and “Out” in the labels denote the embeddings of the second layer and the embeddings of the output, respectively.

Building on these quantitative comparisons (Fig. 8), the 4096-color analysis suggests two distinct geometric structures as testable candidate predictions for the large-scale structure of human color representation. The first candidate is a relatively simple, undistorted ring-like structure, as exemplified by the second-layer embeddings of the three learning paradigms (Fig. 7B-D, H-J, N-P). The consistency observed among the second-layer embeddings of the MAE models (Fig. 8) further suggests that DNNs can converge on a consistent color representation across a vast color space without either linguistic information or supervisory signals. The second candidate is a relatively complex geometry, namely a hollow, distorted ring with specific protrusions and indentations, as exemplified by the output embeddings of the CLIP models (Fig. 7Q-S). This result suggests that the integration of visual information with rich, descriptive language leads to convergence on this distinct geometric structure. Taken as candidate predictions, these two structures define three possibilities for the large-scale structure of human color representation: it may resemble the relatively simple, undistorted ring-like structure exemplified by the second-layer embeddings, the relatively complex geometry exemplified by the output embeddings of the CLIP paradigm, or neither. Determining which of these possibilities corresponds to human color representation remains a question for future large-scale psychophysical experiments.

## Discussion

In this study, we aimed to provide predictions of human color perception across a vast color space by systematically investigating the conditions under which DNNs acquire a color representation that mirrors human color perception. Our results revealed that the early layers of the three learning paradigms and the output of the CLIP paradigm acquire color representations that align with the 93-color human perception (Fig. 5). We found that the second-layer embeddings of the three learning paradigms and the output embeddings of the CLIP paradigm that successfully aligned with human data also developed remarkably consistent and robust color representations in a vast color space (Fig. 7B-D, I, J, O, P, Q-S, Fig 8), allowing us to make novel, testable predictions about the large-scale structure of human color perception. Here, we discuss the implications of these findings with a focus on four key areas: the effect of language on color representations of DNNs, the reason why the output embeddings of the models without rich linguistic input fail to maintain human-like color representations, the use of DNNs as computational proxies for predicting large-scale human color perception, and future prospects for validating and extending these DNN-based predictions.

### Effect of language on color representations of DNNs

The findings that the early layers of the three learning paradigms converged on a relatively undistorted ring-like structure (Fig. 7B-D, H-J, N-P, 8) whereas the output of the CLIP paradigm converged on a ring-like structure with specific protrusions and indentations (Fig. 7Q-S, 8) lead us to consider the possibility that these distortions reflect the influence of language on color representations. We draw this inference because the learning paradigms that align with each other at their early layers include MAE (Fig. 6, 8), which forms representations from visual information alone without any linguistic input or supervisory signals, suggesting that the influence of language on these representations is limited. In contrast, the output of the CLIP paradigm forms representations through the integration of visual information with rich linguistic descriptions, suggesting that the influence of linguistic information is substantial. This language-driven interpretation connects our findings to the long-standing debate on whether language shapes human color perception [72–79], with prior studies addressing this question including investigations using DNNs [42–44, 47, 52]. If the influence suggested by our findings holds, it would lean toward the relativist view that language can shape color representations [72, 74, 75, 77, 79]. However, this raises an important open question: since the CLIP models analyzed in our study were predominantly trained on English-language descriptions, how would their color representations differ if they were trained on other languages? Future studies could address this question by comparing CLIP models trained on diverse languages.

Turning our focus to the early layers, the shared representational structure across the three learning paradigms allows for an intriguing developmental hypothesis: these early representations might correspond to the color perception of pre-linguistic infants. As discussed above, these early layers likely form representations with limited linguistic influence, potentially serving as a precursor to the adult color representation refined by language [80–84]. This linguistically refined representation of adults would correspond to the output of the CLIP paradigm. Under this view, the 93-color human data used in this study [18] may reflect both this fundamental pre-linguistic representation and the subsequent linguistically refined representation of adults. Testing this hypothesis would require comparing the early layers of the three learning paradigms with color representations of pre-linguistic infants or children. However, the detailed representational structure of infants or children in a large color space remains unexplored. Although a recent study reported that children as young as 3 years old already possess a color representation that resembles that of adults in a limited 9-color set [12], this coarse similarity does not guarantee alignment in a larger color space. Accordingly, attempts to determine whether language acquisition is indeed necessary for forming the adult-like color representation require obtaining large-scale color structure data from children or infants and comparing them with the early layers of these learning paradigms. This endeavor would yield significant insights into the developmental trajectory of human color perception.

### Reason why the output embeddings of the models without rich linguistic input fail to maintain human-like color representations

A possible explanation for why models without rich linguistic input fail to maintain human-like color representations at the output (Fig. 5) lies in the known processing dynamics of DNNs. A recent study has demonstrated that color information is prominently represented in early layers of DNNs but progressively diminishes in deeper layers, as the network increasingly prioritizes higher-level, task-specific features such as object categories [37–41, 85]. Our layer-by-layer results (Fig. 5) are consistent with this pattern: models with MAE and SL paradigms achieve strong alignment with human color similarity structure in early or intermediate layers, but this alignment is lost at the output where representations are shaped by their respective training objectives of pixel reconstruction and object classification. In contrast, the CLIP paradigm employs a contrastive objective that aligns the output of the image encoder with that of the language encoder [55], and since linguistic descriptions include human-defined color information, this objective may provide a learning signal that sustains human-like color similarity structure through to the output, even though the deeper layers of the CLIP image encoder have been shown to contain a smaller number of color-selective neurons [40]. This difference in training objectives offers a plausible account of why only CLIP models exhibit human-like color representations at the output, even though all paradigms appear capable of forming such representations at their early layers.

Another possible explanation for why MAE and SL models fail to align with human data at the output (Fig. 5) is their significantly smaller training dataset sizes. Specifically, the volume of the ImageNet dataset [67] falls short of that of the datasets used to train CLIP models [68–71] by more than an order of magnitude. Notably, this size difference is not accompanied by a difference in color, as the hue distribution of ImageNet-1k closely matches that of DFN-2B, the dataset used to train the CLIP models in the main results (Fig. S13). Because scale, unlike color, remains uncontrolled, training MAE and SL models on datasets of a comparable scale would allow for a more rigorous comparison between the learning paradigms. However, due to the massive computational resources required for such a large scale analysis, we leave this investigation for future research.

### DNNs as computational proxies for predicting large-scale human color perception

While perceptually uniform color spaces such as CIELAB [86, 87] have been foundational in color science, previous studies have suggested that they fall short when mapping global relationships across a broader range of colors [88–90], and our findingsare consistent with this view (Fig. S6). Although these classical models were based on a large number of human color judgments, the measured color pairs were typically local, restricted to those that were adjacent [6] or along specific axes [3, 4]. Consequently, these color space models were primarily optimized to capture perceptual color differences between similar colors, ensuring that small Euclidean distances in the space correspond to differences in human color perception. Therefore, there is no guarantee that the distances between colors positioned far apart in these spaces reflect human perceptual dissimilarities [88–90].Our evaluation demonstrated that these color space models did not achieve top-1 matching rates as high as those of DNNs when compared against the human behavioral data capturing global relationships (Fig. S6). However, the known shortcoming of these color spaces for colors positioned far apart is only a potential explanation for their lower top-1 matching rates. Because the top-1 matching rate is determined by the entire similarity structures being compared, it is generally difficult to identify which local part of the structures causes a low top-1 matching rate.Nevertheless, the lower top-1 matching rates indicate thatthese color space models have limitations in exploring the large-scale, global structure of human color perception. Furthermore, modifying these spaces to achieve global uniformity via traditional psychophysical methods is impractical due to the combinatorial explosion of required measurements.

To overcome the methodological bottleneck of exhaustively measuring an enormous number of color combinations, human-aligned DNNs could offer a powerful computational approach. In contrast to these color space models, some DNN representations showed strong alignment with the human behavioral data capturing global color relationships (Fig. 5). By utilizing these human-aligned models as computational proxies, we generated testable predictions of the perceptual maps of a vastly expanded space comprising 4,096 colors (Fig. 7B-D, I, J, O, P, Q-S). Evaluating these DNN-generated large-scale predictions against actual human data could help address the outstanding issues of global non-uniformity in current models and guide the development of globally robust perceptually uniform color spaces.

### Future prospects for predicting human color perception with DNNs

The first future prospect involves empirically testing the plausibility of our prediction for the vast, 4096-color representation suggested by the human-aligned DNN models (Fig. 7 B-D, I, J, O, P, Q-S). In order to validate, we naturally need to compare our results with human data. This would require psychophysical experiments comparing the vast number of color pairs necessary to construct the 4096-color representation, which is infeasible for an individual subject to conduct. Therefore, one potential approach is to collect data from a group of people, as utilized in the previous research [18]. Although computationally and logistically demanding, empirical validation of this large-scale prediction remains a critical next step for future research.

The second important future direction is to investigate more naturalistic and nuanced color representations. A limitation of the current study, consistent with much foundational research in this area, is the use of idealized, single-color patches. This controlled approach is necessary for isolating variables, but does not capture the complexities of color perception in natural environments. Human color perception is heavily influenced by context, such as changing lighting conditions [91], the influence of surrounding backgrounds [92], and the complex interplay of multiple colors [93]. In our own analyses, the alignment with human data already changed when the color patches were surrounded by a grey background (Fig. S3). Given this finding, and because our results rely on simple stimuli, it remains unclear how the color representations reported here hold for colors embedded in richer, more naturalistic stimuli. Future work should therefore extend this research to examine whether the human-like alignment of DNN models holds under these more realistic conditions.

## Materials and methods

### Human Behavioral Data

Human behavioral data were drawn from a large-scale dataset collected from 426 color-neurotypical participants in a previous study [18]. In the experiment, participants performed a color similarity judgment task in which they were asked to rate the perceived similarity between pairs of 93 color stimuli selected from the Practical Color Coordinate System (PCCS) [66]. For each pair, ratings were provided on a scale from 0 (“very similar”) to 7 (“very different”). The human dissimilarity matrix was computed as the pairwise Euclidean distances between 20-dimensional color embeddings estimated from the participants’ similarity judgments [18].

### DNN Models Used in This Study

We used three types of deep neural network (DNN) learning methods in this study: self-supervised learning, supervised learning, and CLIP (Contrastive Language–Image Pre-training). For each of these learning paradigms, we used publicly available pretrained models based on the Transformer architectures ViT-B, ViT-L, and ViT-H and the CNN architectures ResNet-50 and ResNet-101. We used the same architectures across the three learning paradigms so that the differences we observed would reflect the learning paradigm rather than the architecture. We also included CNN-based architectures in addition to Transformer-based ones so that our findings would not be specific to a single class of architectures. A brief description of each learning paradigm is provided below.

### Self-Supervised Learning models

Self-supervised learning (SSL) is a framework where models learn representations from raw data by solving pretext tasks without relying on supervised signals. For our study investigating color representation, it was crucial to select models that had not been trained with color augmentation, as such techniques could artificially degrade or alter the intrinsic color information learned by the model. Our selection was therefore guided by this constraint, and lead us to focus on masked image modeling (MIM) models [53, 54, 94]. In this framework, the model learns by masking random patches of the input natural image and reconstructing the missing raw pixel values. These models learned representations exclusively from the input images themselves, without any reliance on labels or linguistic information.

### Supervised Learning models

Supervised learning (SL) is a paradigm where models learn to classify or predict outputs from the input natural image [22, 25]. These models were trained on labeled datasets of natural images. Although the labels were not primarily focused on color information, some inherently included color descriptors (e.g., “black swan”, “red wine”), indirectly providing the models with color-related information during training.

### CLIP models

CLIP (Contrastive Language–Image Pretraining) models learn to associate natural images and text by jointly training on large-scale image-text pairs [55]. The architecture consists of separate image and text encoders that project visual and linguistic features into a shared embedding space, where the model is trained to maximize the similarity of matched pairs via a contrastive objective. Consistent with the other learning paradigms, we used only the image encoders for our analysis. These models were trained with free-form natural language descriptions that objectively describe the visual content of an image, typically as a single short sentence or phrase, and these descriptions included color descriptions. Unlike curated class labels, color in these descriptions is addressed naturally as a visual attribute modifying a specific object (e.g., “A green highway sign with the words Queens Bronx.”). These natural language descriptions enable the models to learn representations that link visual features with color-related linguistic information.

### Training Datasets of the DNN Models

The DNN models were trained on different datasets depending on their learning paradigm: ImageNet-1k for the MAE and SL models, and substantially larger image-text datasets for the CLIP models. ImageNet-1k [67] consists of *∼*1.28 million object-centric natural images that were collected from the Internet and labeled by human annotators. The datasets used to train the CLIP models consist of natural images paired with text descriptions drawn from the web, with the image-text pairs selected by automatic filtering. These were YFCC [68] for the CNN-based models, and LAION-2B [69], MetaCLIP [70], or DFN [71] for the Transformer-based models, together comprising *∼*15 million to *∼*5 billion image-text pairs.

Despite these differences in scale and assembly, the datasets are comparable in their visual and color content, with no curation toward particular colors. Both span a similar breadth of everyday content, such as objects, animals, people, scenes, and architecture. To assess their color content directly, we compared the hue distribution of ImageNet-1k with that of DFN-2B (the dataset used to train the CLIP models in the main results) following the method of [37], and found the two distributions to be very similar (Fig. S13). The shape of these distributions, in which the orange region contains the largest share of chromatic samples followed by a secondary peak on the blue side, is consistent with the color statistics reported across a wide range of image and reflectance databases [95].

### Deriving Representational Dissimilarity Matrices from DNNs

To create model-based dissimilarity matrices that were comparable to the human behavioral data, we performed the following procedure for each DNN model:

1. We generated 93 images corresponding to the exact same PCCS color stimuli used in the human behavioral experiment [18]. For the main analysis, each image was uniformly filled with one of the 93 colors. For the analysis presented in the Supporting information (Fig. S3), we compared the representations of these uniformly colored images with those of images containing a central circular patch for each of the 93 colors on a grey (#7F7F7F) background. The radii of the circular patches were set to 1/2, 1/3, and 1/4 of the image width.

2. Each image was individually fed into a model, from which we extracted the corresponding feature embedding from the image encoder as follows.

• Transformer-based models: The CLS token at each layer and the output of its image encoder was extracted as the embedding. For object classification models, we treated the final layer embedding with layer normalization applied as the output embedding.

• CNN-based models: Embeddings were extracted from the initial max-pooling layer, the first two Bottleneck blocks of each of the four residual stages, and the output of its image encoder. At each layer, the output of that layer, which is a feature map, was flattened into a one-dimensional vector, and this vector was used as the embedding. For object classification models, we treated the output of the fourth residual stage with global average pooling applied as the output embedding.

3. We then computed a 93 × 93 representational dissimilarity matrix (RDM) by calculating the pairwise Euclidean distances between the extracted 93 color embeddings.

The same procedure was also applied to derive RDMs for two larger color sets. The first was a richer 297-color set from the Practical Color Coordinate System (PCCS), comprising 288 chromatic colors (24 hues × 12 tones) and 9 achromatic colors. The second was a 4096-color set designed to exhaustively cover the entire digital RGB space, generated by uniformly sampling 16 values from each of the R, G, and B channels (16^3^ = 4096). The findings from the 297-color set were highly consistent with the 93-color experiment. As this analysis confirmed our main results rather than uncovering novel large-scale structures as the 4096-color experiment did, we include it in the supporting information to support our primary conclusions (Fig. S12).

### RGB color space baseline

To establish a simple, non-learned baseline for comparison, we computed a representational dissimilarity matrix (RDM) for the RGB color space. For each color set (93 PCCS colors, extended 297 PCCS colors, and 4096 RGB colors), the RDM was computed by calculating the Euclidean distance between each color pair based on their coordinates in the 3-dimensional RGB space.

### Color space models other than RGB

To provide additional baselines that incorporate established knowledge of human color vision, we computed RDMs for four more advanced color spaces: the cone-opponent DKL space, the perceptually uniform *J_z_a_z_b_z_* and CIELAB spaces, and the CIECAM16 color appearance model. For each of these spaces, the 93 PCCS colors were converted from their sRGB values into the corresponding coordinates, and the 93 × 93 RDM was computed as the pairwise Euclidean distances between these coordinates. All conversions were performed with the colour-science Python library (version 0.4.7) [96], treating the input colors as sRGB values under a D65 white point. Each color space and its conversion are described as follows:

- DKL (Derrington–Krauskopf–Lennie): a physiologically motivated, cone-opponent color space [97]. The sRGB values were converted to CIE 1931 XYZ tristimulus values and then to LMS cone excitations based on the Stockman & Sharpe 2*^◦^* cone fundamentals [98]. Each color was expressed as Weber cone contrasts relative to a D65 adapting white, and these contrasts were projected onto the three cardinal DKL axes: an achromatic luminance axis (*L* + *M* ), a red–green axis (*L − M* ), and a blue–yellow axis (*S −* (*L* + *M* )).
- *J_z_a_z_b_z_*: a perceptually uniform color space designed for high-dynamic-range and wide-gamut image signals [99]. The sRGB values were converted to CIE 1931 XYZ and then to *J_z_a_z_b_z_* coordinates—a lightness axis *J_z_* and two chromatic axes *a_z_* and *b_z_*—under the D65 white point. The Euclidean distance between two colors in this space approximates the perceived color difference.
- CIELAB: a perceptually uniform color space standardized by the CIE [86]. The sRGB values were converted to CIE 1931 XYZ and then to CIE *L^∗^a^∗^b^∗^* coordinates under the D65 white point and the 2*^◦^* standard observer. In this space, the Euclidean distance between two colors corresponds to the CIE76 color difference 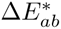 .
- CIECAM16 (CAM16-UCS): a color appearance model that predicts perceived color attributes under specified viewing conditions [100]. From the XYZ values we computed the CIECAM16 appearance correlates—lightness (*J*), colorfulness (*M* ), and hue angle (*h*)—under a D65 white point, an average surround, an adapting field luminance of *L_A_* = 16 cd*/*m^2^, and a relative background luminance of *Y_b_* = 20. To obtain a space in which Euclidean distance approximates the perceived color difference, these correlates were transformed into the uniform CAM16-UCS coordinates (*J′*, *a′*, *b′*) [100].

### Alignment of the MDS configurations to the human representation

To make the MDS configurations directly comparable across models, we aligned the orientation of every two- and three-dimensional MDS configuration of the models in this study to that of the human 93-color configuration of the same dimensionality (Fig. 2F and Video S1). Because an MDS configuration is determined only up to rotation and reflection, the orientation in which it is displayed is arbitrary, and changing this orientation does not alter the representation itself. For each model’s configuration, we obtained the orthogonal Procrustes transformation that minimized the distances between the positions of the colors used in the human experiment in the model’s configuration and those of the same colors in the human configuration, and applied this transformation to all colors in the model’s configuration. Because this transformation is restricted to rotation and reflection and involves neither scaling nor shearing, it changes only the orientation of a configuration and leaves all distances within it unchanged.

This alignment required the positions of the colors used in the human experiment in each model’s configuration, and how we obtained these positions depended on which colors the configuration contained. The 93-color configurations (Fig. 2G-J, S2B-K, S6H-L, S9B-K, and Video S1–S3) contain all 93 of these colors, and we used their positions. The 297-color configurations (Fig. S12A-J) contain 89 of the 93 colors, and we used the positions of these 89 colors. The 4096-color configurations (Fig. 7 and Video S4–S6) contain none of the 93 colors, and we therefore placed the 93 colors into each of these configurations on the basis of their distances to the 4096 colors. For each model and layer, we computed the Euclidean distances between the embedding of each of the 93 colors and the embeddings of the 4096 colors in the same embedding space, and divided them by the maximum distance among the 4096 colors, the same value by which the dissimilarity matrix of the 4096 colors had been normalized. We then determined the position of each of the 93 colors by minimizing the stress between these normalized distances and the distances from the position to the 4096 points of the configuration.

### Supervised comparison based on Pearson correlation

To quantify the similarity between any two representational dissimilarity matrices (RDMs) in a supervised manner, we computed the Pearson correlation coefficient (*ρ*), known as conventional Representational Similarity Analysis (RSA) [56–58]. The correlation is calculated as:

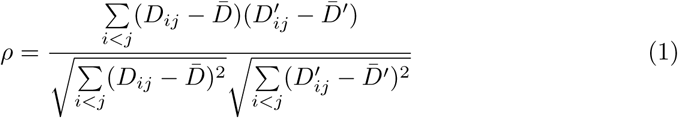

where *D_ij_* and 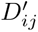 are the dissimilarities between color *i* and color *j* in the first RDM (*D*) and the second RDM (*D′*), respectively. The terms *D̄* and *D̄′* represent the mean of all dissimilarity values in the respective RDMs. The summation is performed over the upper triangular elements of the matrices (where *i < j*) to include each color pair only once. The central assumption of this method is that a known one-to-one correspondence exists between the colors of the two RDMs, which makes a direct comparison between the dissimilarity of “color A vs. color B” in one RDM and “color A vs. color B” in the other a meaningful measure of similarity.

### Unsupervised comparison based on Gromov–Wasserstein Optimal Transport (GWOT)

#### Gromov–Wasserstein Optimal Transport (GWOT)

We employed the Gromov–Wasserstein optimal transport (GWOT) algorithm [60, 61] for the unsupervised comparison of color similarity structures. This approach is used to assess structural similarity without relying on any pre-defined correspondence between the colors in the different structures. GWOT functions as an unsupervised alignment method that determines an optimal transportation plan between two distinct point clouds, and operates without any item-specific correspondence data. The core of this method involves the optimization (minimization) of the Gromov–Wasserstein distance (GWD), defined as:

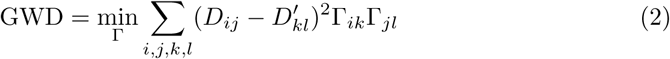

GWD serves as a metric to quantify the level of correspondence between the two similarity structures (Fig. 9A). For this study, *D_ij_* denotes the dissimilarity between colors *i* and *j* in the first similarity matrix *D*, and *D′* denotes the dissimilarity between colors *k* and *l* in the second matrix *D′*. Prior to computation, every similarity matrix was normalized so that its values fell within a [0, 1] range. The solution to this GWD minimization problem is the optimal transportation plan, denoted as the matrix Γ. This matrix Γ provides an effective unsupervised alignment of the two color similarity structures. Each element Γ*_ik_* within this matrix can be understood as the “probability” that the *i*-th color from the first domain maps to the *k*-th color in the second domain.

**Fig 9.**
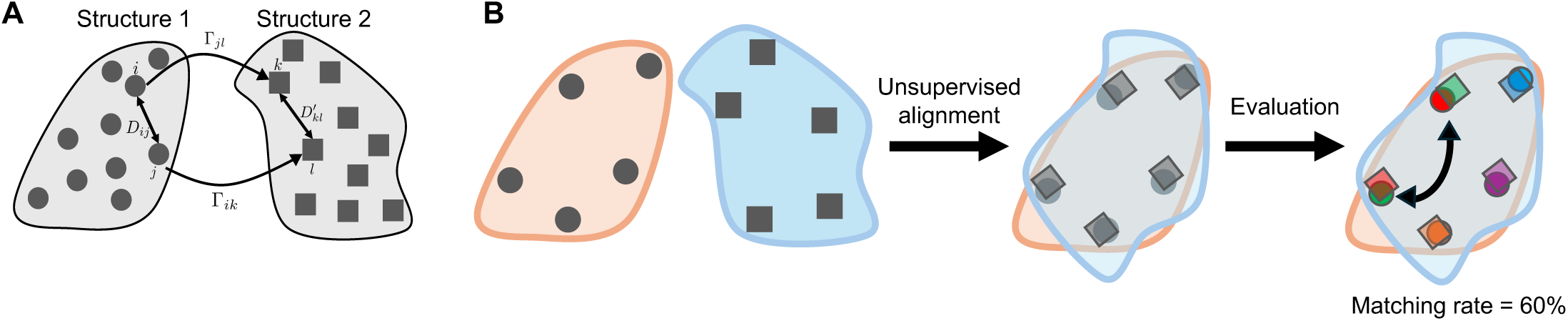
Schematic of the unsupervised comparison based on Gromov–Wasserstein Optimal Transport (GWOT). (A) Schematic of Gromov–Wasserstein optimal transport. The elements of matrices *D* and *D′* are the dissimilarities between the items. Γ is the transportation matrix, where each element indicates the probability of an item in one similarity structure corresponding to another item in the other similarity structure. (B) Evaluation of unsupervised alignment using external labels. Following unsupervised alignment based solely on the internal dissimilarities of each structure, the matching rate between them was calculated using ground-truth labels.

Optimizing the GWD (Eq. 2) presents a non-convex problem. Consequently, any solution found by the algorithm is a local optimum, and there is no assurance of reaching the global optimum. To manage this optimization challenge and identify good local minima, we employed the Python Optimal Transport (POT) library [101]. We executed the algorithm multiple times, each time beginning with a different random initialization of the Γ matrix. Specifically, 5 × 10^6^ random initializations were performed for the 93-color dissimilarity matrices in the analysis of the output embeddings, 5 × 10^5^ random initializations were performed for the 93-color dissimilarity matrices in the layer-by-layer analysis, and 1 × 10^5^ initializations were performed for the 297-color matrices.

An alternative and more efficient method for optimizing GWD involves the inclusion of an entropy-regularization term, *H*(Γ). This modification results in the following objective function:

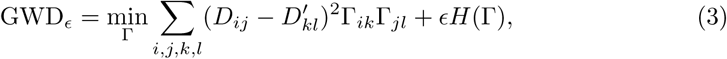

The introduction of this term is known to improve optimization efficiency [59, 102]. However, this regularized problem remains non-convex, and accordingly still requires multiple initializations to locate good local minima. We applied this entropy-regularized method to the 4096-color dissimilarity matrices, for which we performed 100 random initializations.

### Quantifying Alignment with Matching Accuracy

To evaluate the quality of the unsupervised alignment produced by GWOT, we calculated a matching rate as the score of matching accuracy. This score measures the extent to which the optimal transportation plan Γ correctly maps corresponding colors between the two structures, using their ground-truth labels for verification(Fig. 9B). Specifically, for each color *i* in the first structure, we identify the color *j* in the second structure to which it is mapped with the highest probability. We consider this a “match” if the label of color *i* is identical to the label of color *j*.

This matching condition can be formalized as follows. Let the color labels for the items in the two dissimilarity matrices be *c*_1_ and *c*_2_. The match for the *i*-th color in the first matrix (*c*_1*i*_) is determined by:

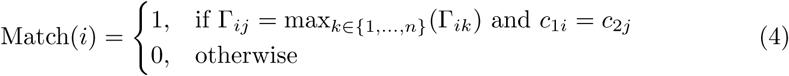

The overall matching accuracy is then the percentage of correctly matched colors, calculated as:

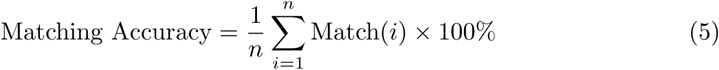

## Supporting information

**Table S1.**
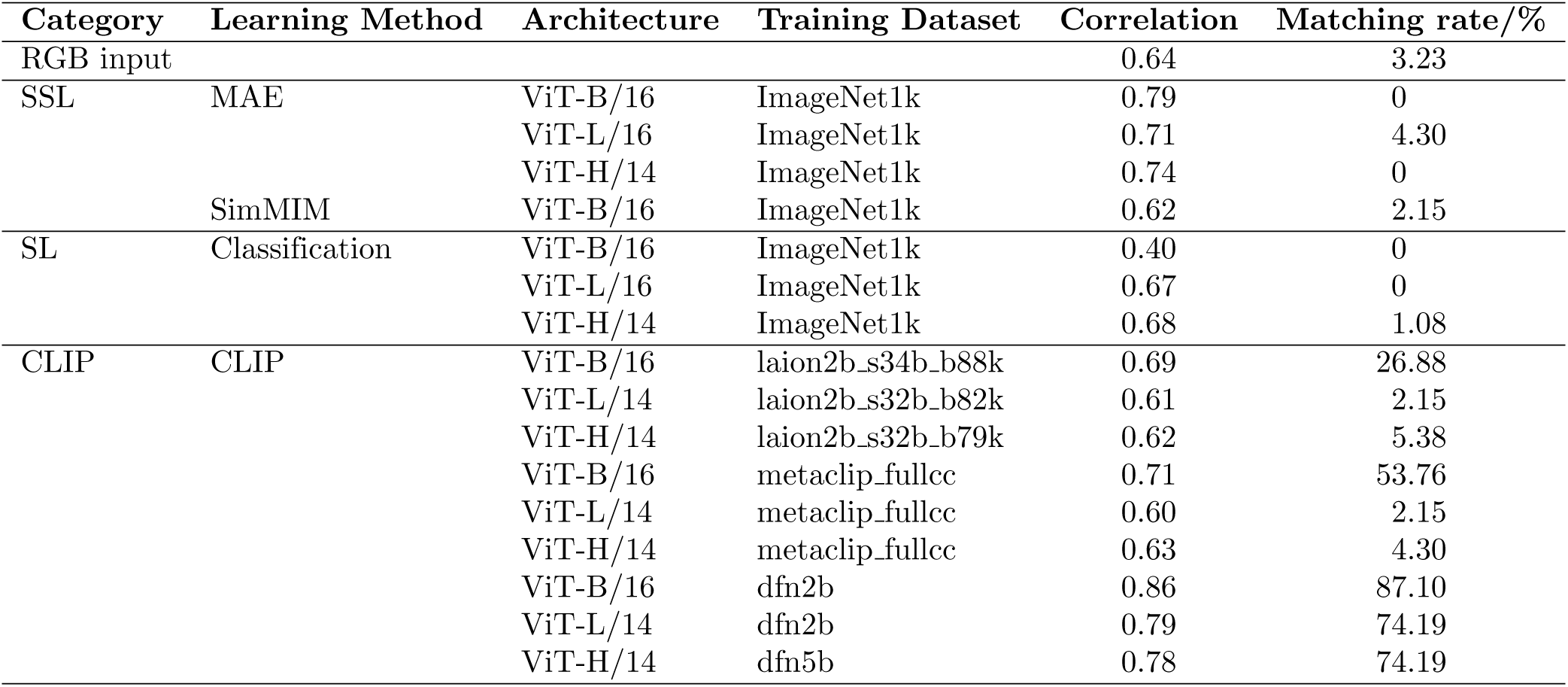
List of the Transformer-based DNN models used in this study and their correspondence of the output embeddings with human data for 93 colors, with 5 × 10^6^ initializations of GWOT.

| Category | Learning Method | Architecture | Training Dataset | Correlation | Matching rate/% |
| --- | --- | --- | --- | --- | --- |
| RGB input |  |  |  | 0.64 | 3.23 |
| SSL | MAE | ViT-B/16 | ImageNet1k | 0.79 | 0 |
|  |  | ViT-L/16 | ImageNet1k | 0.71 | 4.30 |
|  |  | ViT-H/14 | ImageNet1k | 0.74 | 0 |
|  | SimMIM | ViT-B/16 | ImageNet1k | 0.62 | 2.15 |
| SL | Classification | ViT-B/16 | ImageNet1k | 0.40 | 0 |
|  |  | ViT-L/16 | ImageNet1k | 0.67 | 0 |
|  |  | ViT-H/14 | ImageNet1k | 0.68 | 1.08 |
| CLIP | CLIP | ViT-B/16 | laion2b_s34b_b88k | 0.69 | 26.88 |
|  |  | ViT-L/14 | laion2b_s32b_b82k | 0.61 | 2.15 |
|  |  | ViT-H/14 | laion2b_s32b_b79k | 0.62 | 5.38 |
|  |  | ViT-B/16 | metaclip_fullcc | 0.71 | 53.76 |
|  |  | ViT-L/14 | metaclip_fullcc | 0.60 | 2.15 |
|  |  | ViT-H/14 | metaclip_fullcc | 0.63 | 4.30 |
|  |  | ViT-B/16 | dfn2b | 0.86 | 87.10 |
|  |  | ViT-L/14 | dfn2b | 0.79 | 74.19 |
|  |  | ViT-H/14 | dfn5b | 0.78 | 74.19 |

**Fig S1.**
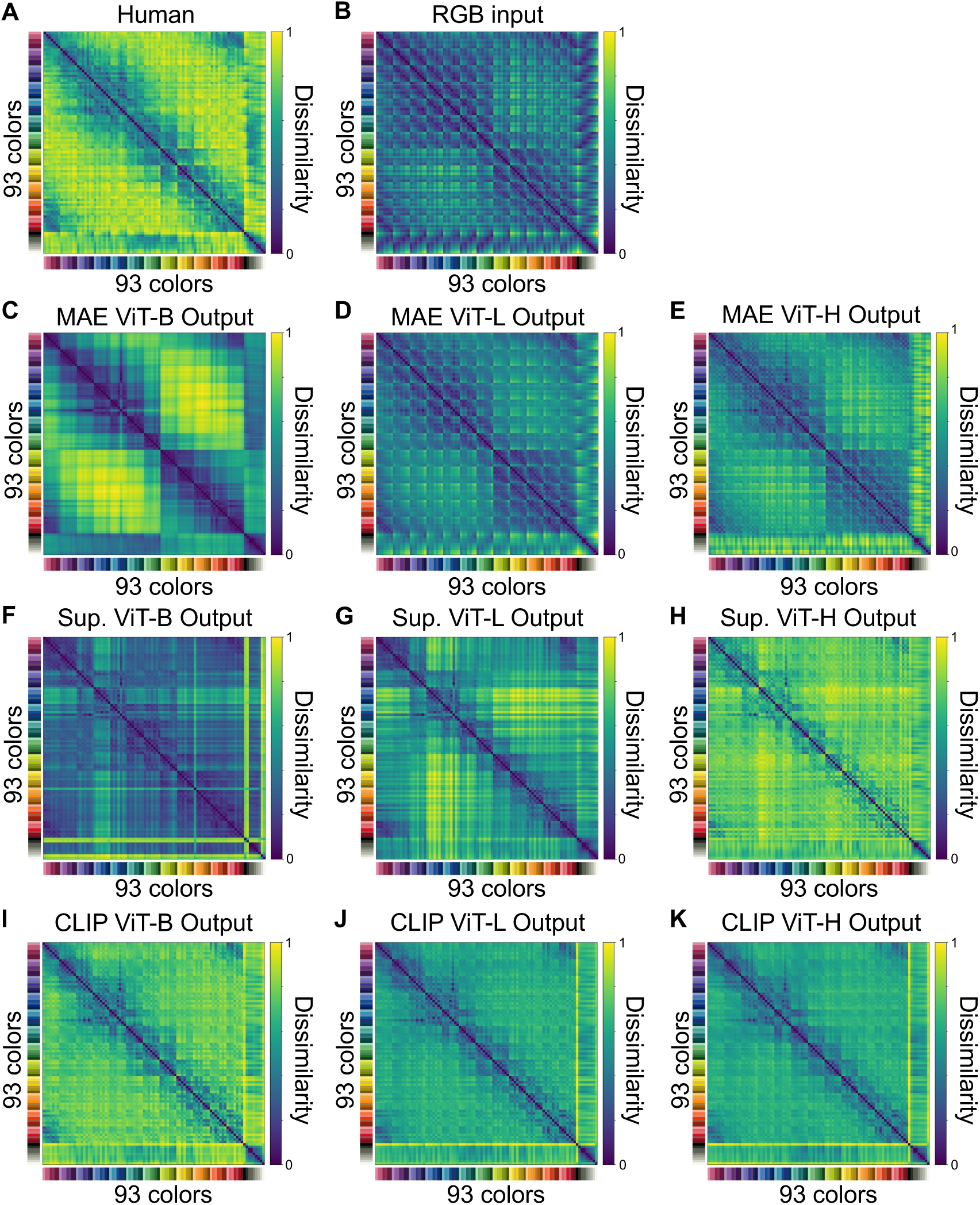
Dissimilarity matrices of 93 colors from humans, RGB input, and the output embeddings of the DNN models. (A)-(K) Dissimilarity matrices of 93 colors from (A) the color-neurotypical participants group, (B) the RGB model, (C)-(E) the MAE models, (F)-(H) the SL models, and (I)-(K) the CLIP models.

**Fig S2.**
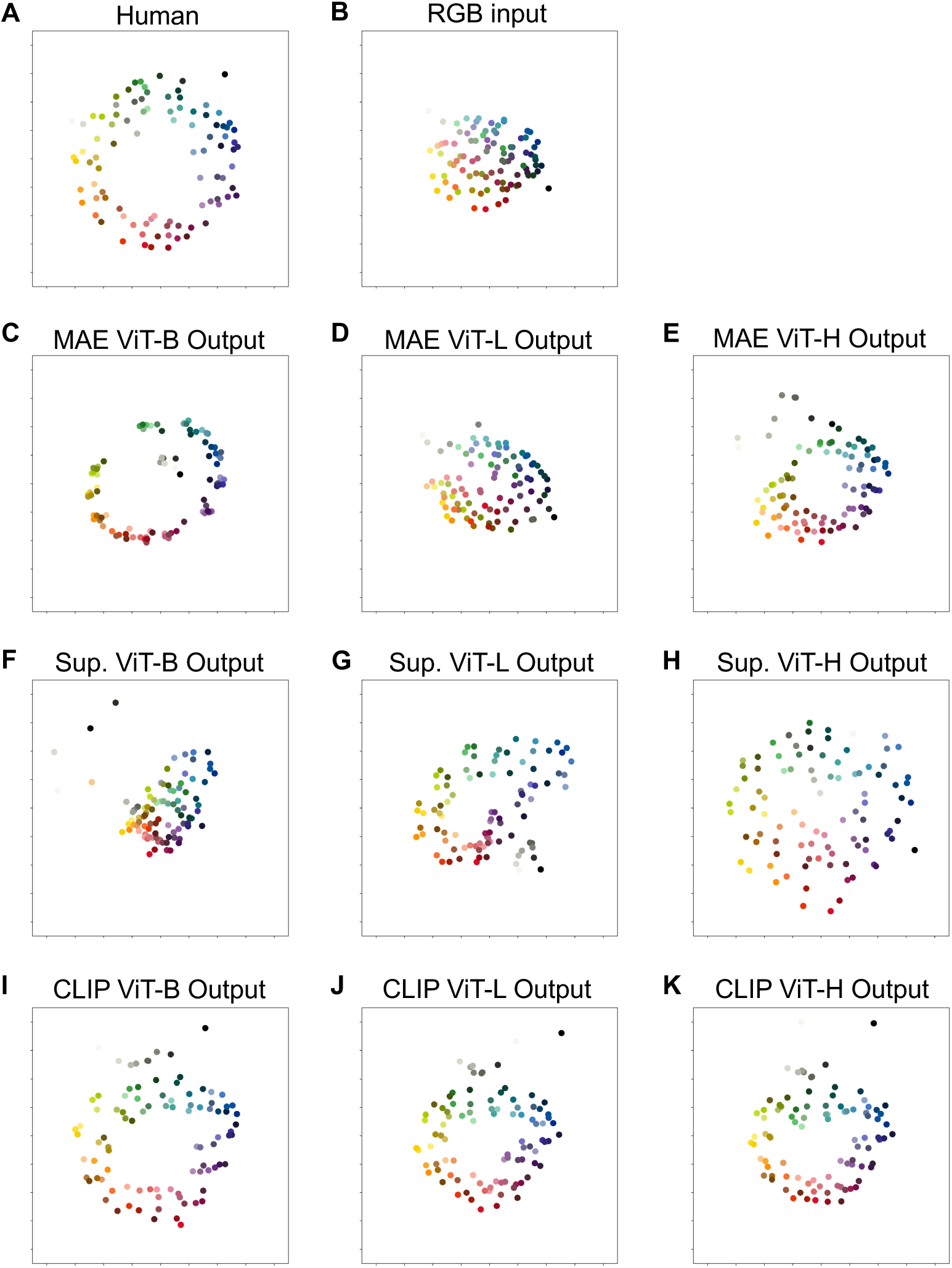
Embeddings of 93 colors from humans, RGB input, and the output embeddings of the DNN models. (A)-(K) Visualization of the embeddings of 93 colors by applying 2-dimensional MDS on the dissimilarity matrix from (A) the color-neurotypical participants group, (B) the RGB model, (C)-(E) the MAE models, (F)-(H) the SL models, and (I)-(K) the CLIP models.

**Fig S3.**
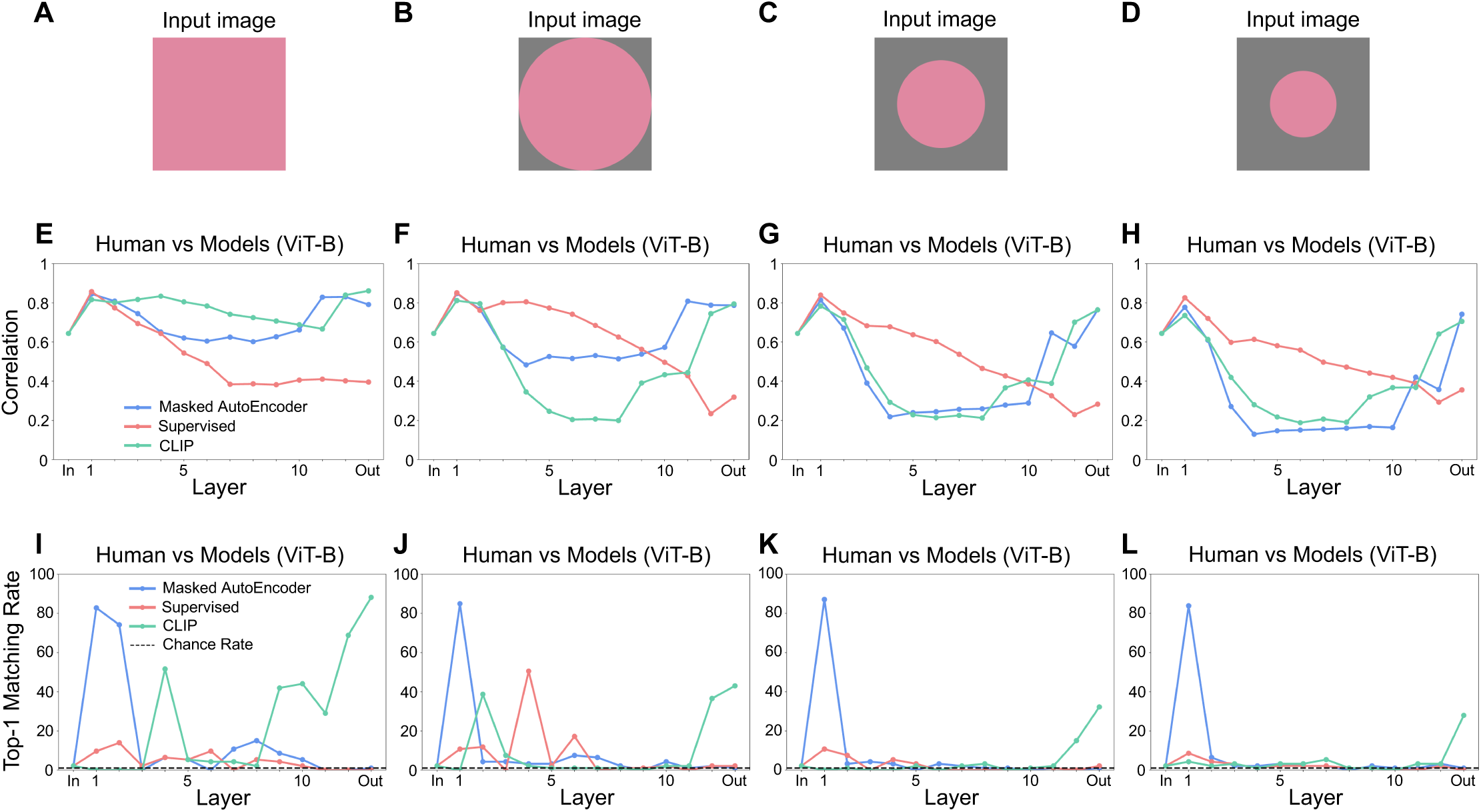
Supervised and unsupervised alignment of color similarity structure for 93 colors between humans and ViT-B-based DNN models across different types of input images. (A)-(D) Sample input images with (A) monotone color patches, (B) circular color patches with a radius equal to 1/2 of the image width on a grey background, (C) circular color patches with a radius equal to 1/3 of the image width on a grey background, and (D) circular color patches with a radius equal to 1/4 of the image width on a grey background. (E)-(H) Supervised comparison based on Pearson correlation between the dissimilarity matrices of humans and the embeddings of each layer of the ViT-B-based DNN models, corresponding to the input images shown in (A)-(D), respectively. (I)-(L) Unsupervised comparison based on GWOT between the dissimilarity matrices of humans and the embeddings of each layer of the ViT-B-based DNN models, corresponding to the input images shown in (A)-(D), respectively. The MAE, SL, and CLIP models shown here are the same models as those presented in the main results: the MAE and SL models trained on ImageNet-1k and the CLIP model trained on DFN-2B. In all panels, “In” and “Out” on the horizontal axis denote the RGB input and the output embedding, respectively.

**Fig S4.**
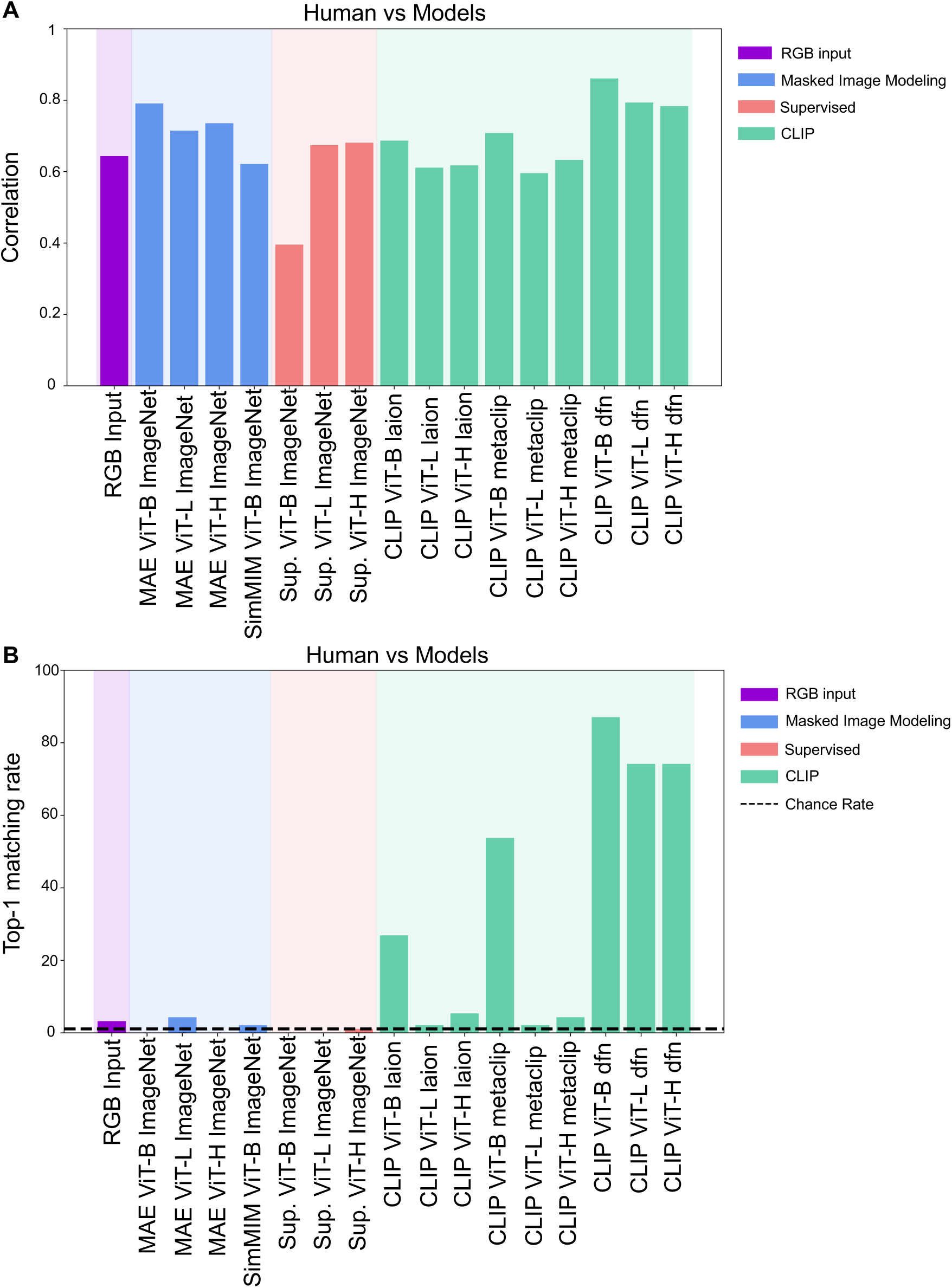
Supervised and unsupervised comparison of color similarity structure for 93 colors between humans and all of the output embeddings of Transformer-based DNN models, with 5 × 10^6^ initializations of GWOT. (A) Supervised comparison based on Pearson correlation. (B) Unsupervised comparison based on GWOT.

**Fig S5.**
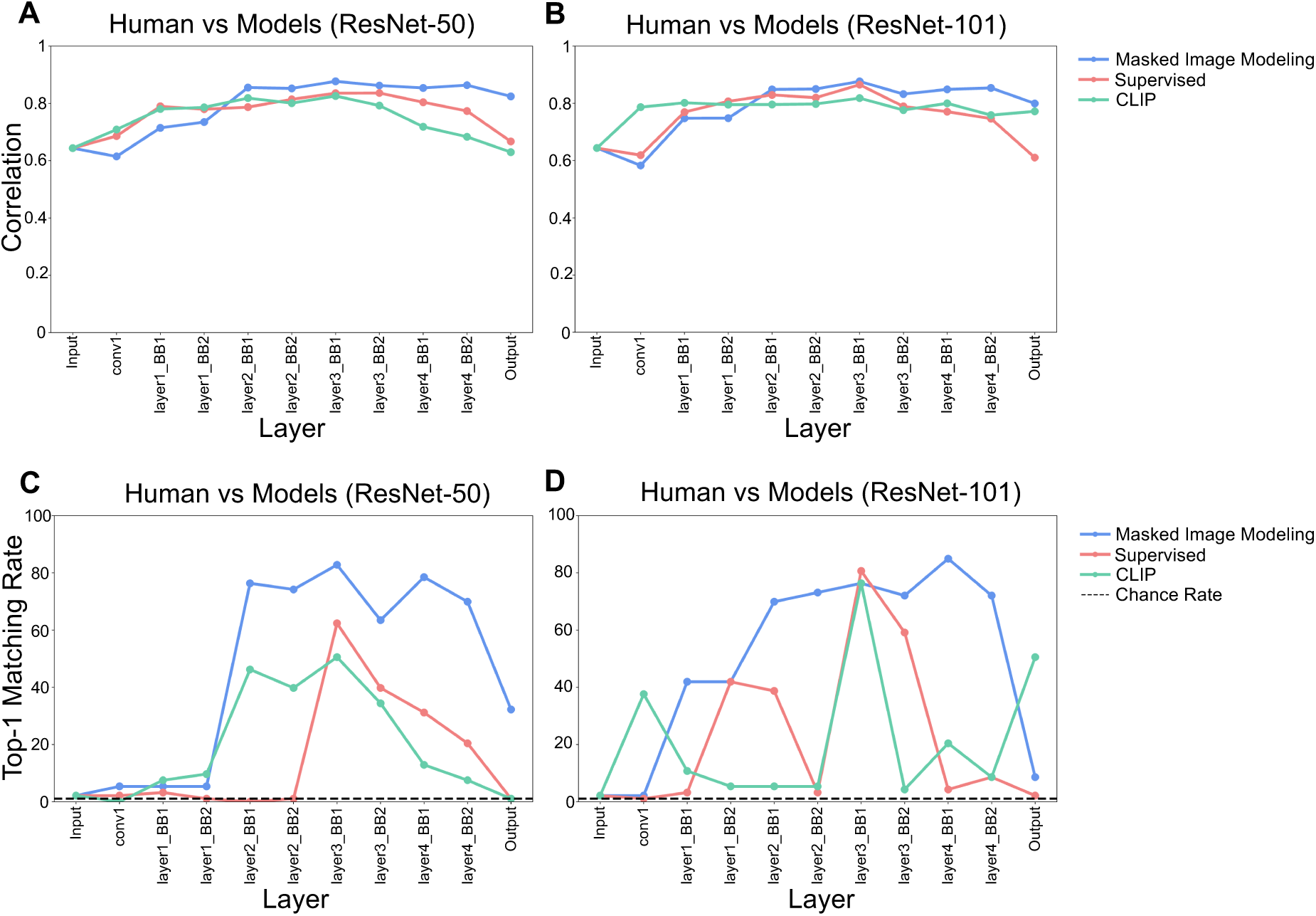
Supervised and unsupervised alignment of color similarity structure for 93 colors between humans and CNN-based DNN models. (A), (B) Supervised comparison based on Pearson correlation between the dissimilarity matrices of humans and the embeddings of each layer of DNN models for (A) ResNet-50 and (B) ResNet-101 architectures. (C), (D) Unsupervised comparison based on GWOT between the dissimilarity matrices of humans and the embeddings of each layer of DNN models for (C) ResNet-50 and (D) ResNet-101 architectures. In all panels, “BB” on the horizontal axis denotes the Bottleneck Block.

**Fig S6.**
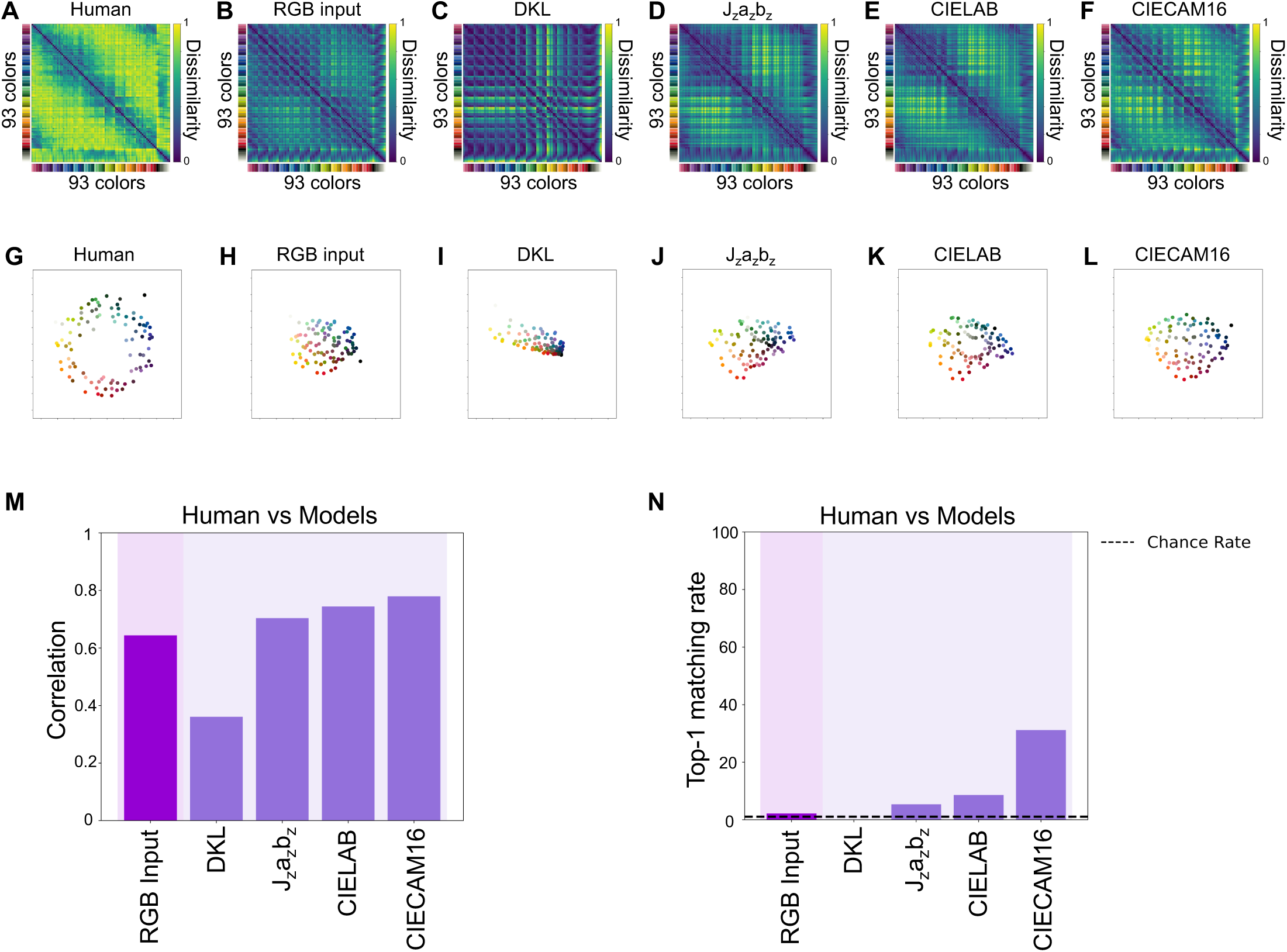
Qualitative and quantitative analysis on color similarity structure for 93 colors between humans, RGB-input and other color space models. (A)-(F) Dissimilarity matrices of 93 colors from (A) the color-neurotypical participants group, (B) the RGB input, (C) DKL, (D) *J_z_a_z_b_z_*, (E) CIELAB, and (F) CIECAM16. (G)-(L) Visualization of the embeddings of 93 colors by applying 2-dimensional MDS on the dissimilarity matrix from (G) the color-neurotypical participants group, (H) the RGB input, (I) DKL, (J) *J_z_a_z_b_z_*, (K) CIELAB, and (L) CIECAM16. (M) Supervised comparison based on Pearson correlation between each pair of humans, RGB input and the other color space models. (N) Unsupervised comparison based on GWOT between each pair of humans, RGB input and the other color space models.

**Fig S7.**
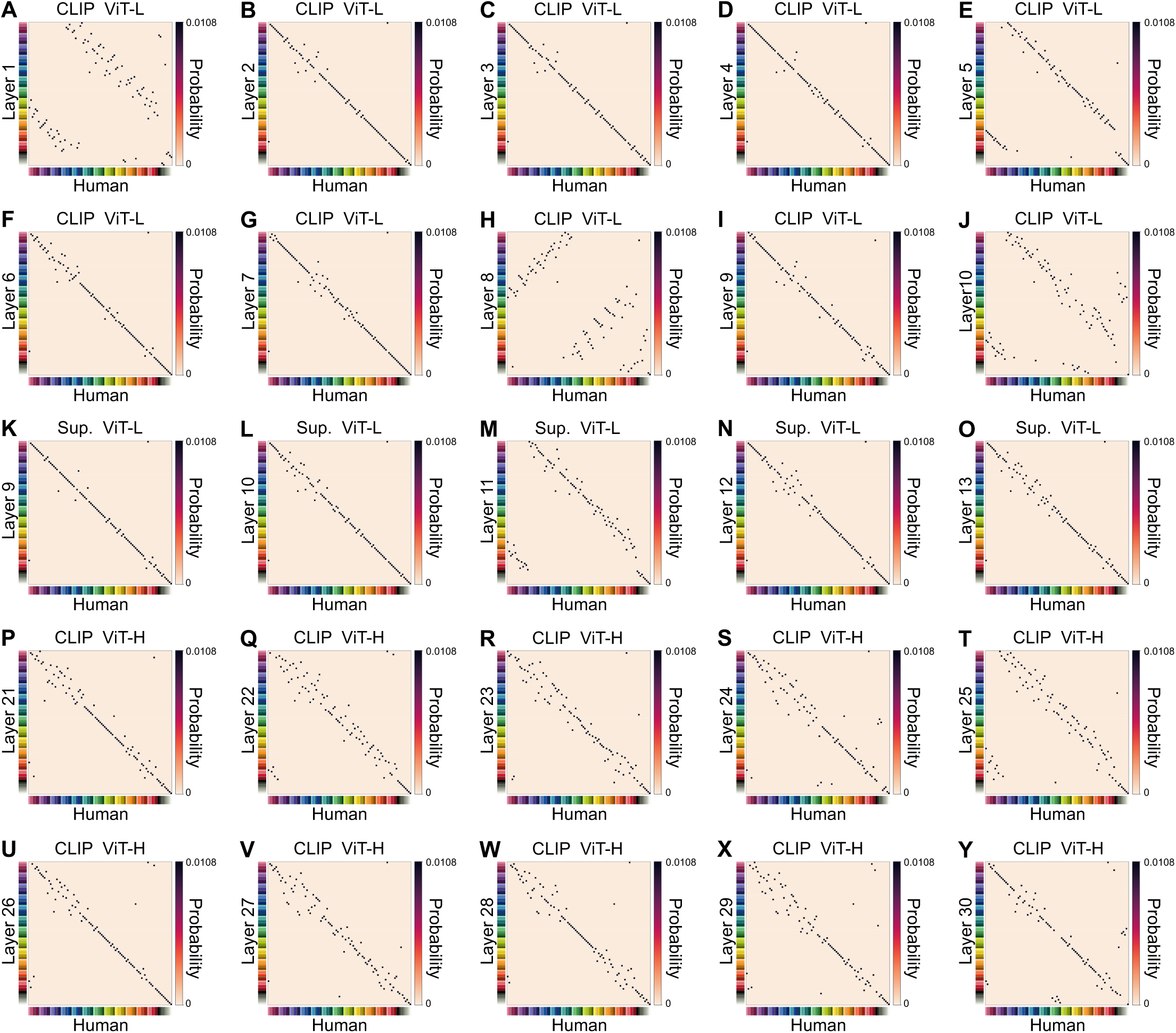
Optimal transportation plans for 93 colors between humans and the embeddings of the layers of DNN models over which the top-1 matching rate fluctuates. (A)-(Y) Optimal transportation plans between the dissimilarity matrices of humans and the embeddings of (A)-(J) the first to tenth layers of the CLIP model in ViT-L, (K)-(O) the ninth to thirteenth layers of the SL model in ViT-L, and (P)-(Y) the twenty-first to thirtieth layers of the CLIP model in ViT-H. In each panel, the horizontal and vertical axes indicate the 93 colors of humans and of the model, respectively, arranged in the same order, so that the diagonal corresponds to the correct one-to-one correspondence.

**Fig S8.**
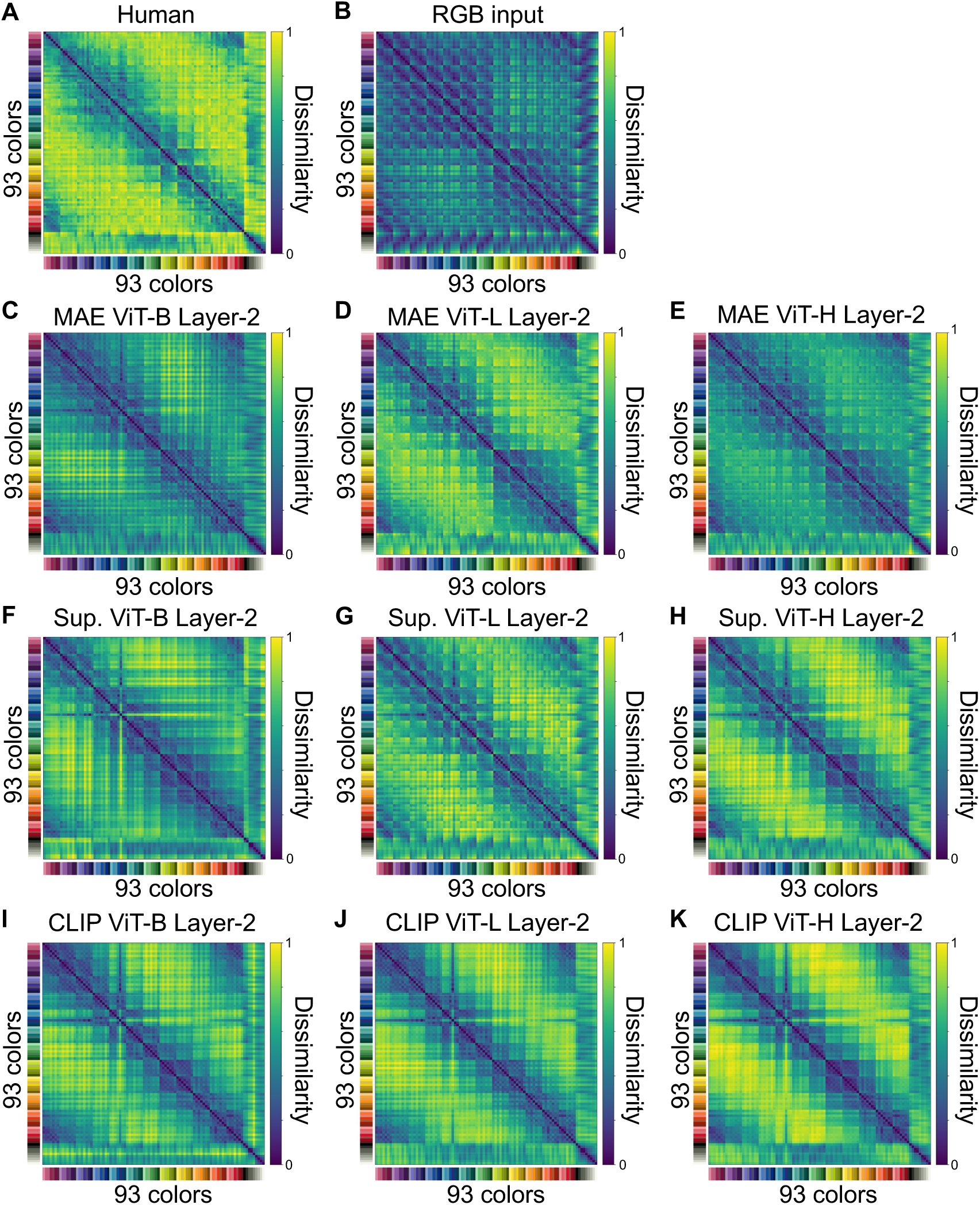
Dissimilarity matrices of 93 colors from humans, RGB input, and the second-layer embeddings of the DNN models. (A)-(K) Dissimilarity matrices of 93 colors from (A) the color-neurotypical participants group, (B) the RGB model, (C)-(E) the MAE models, (F)-(H) the SL models, and (I)-(K) the CLIP models.

**Fig S9.**
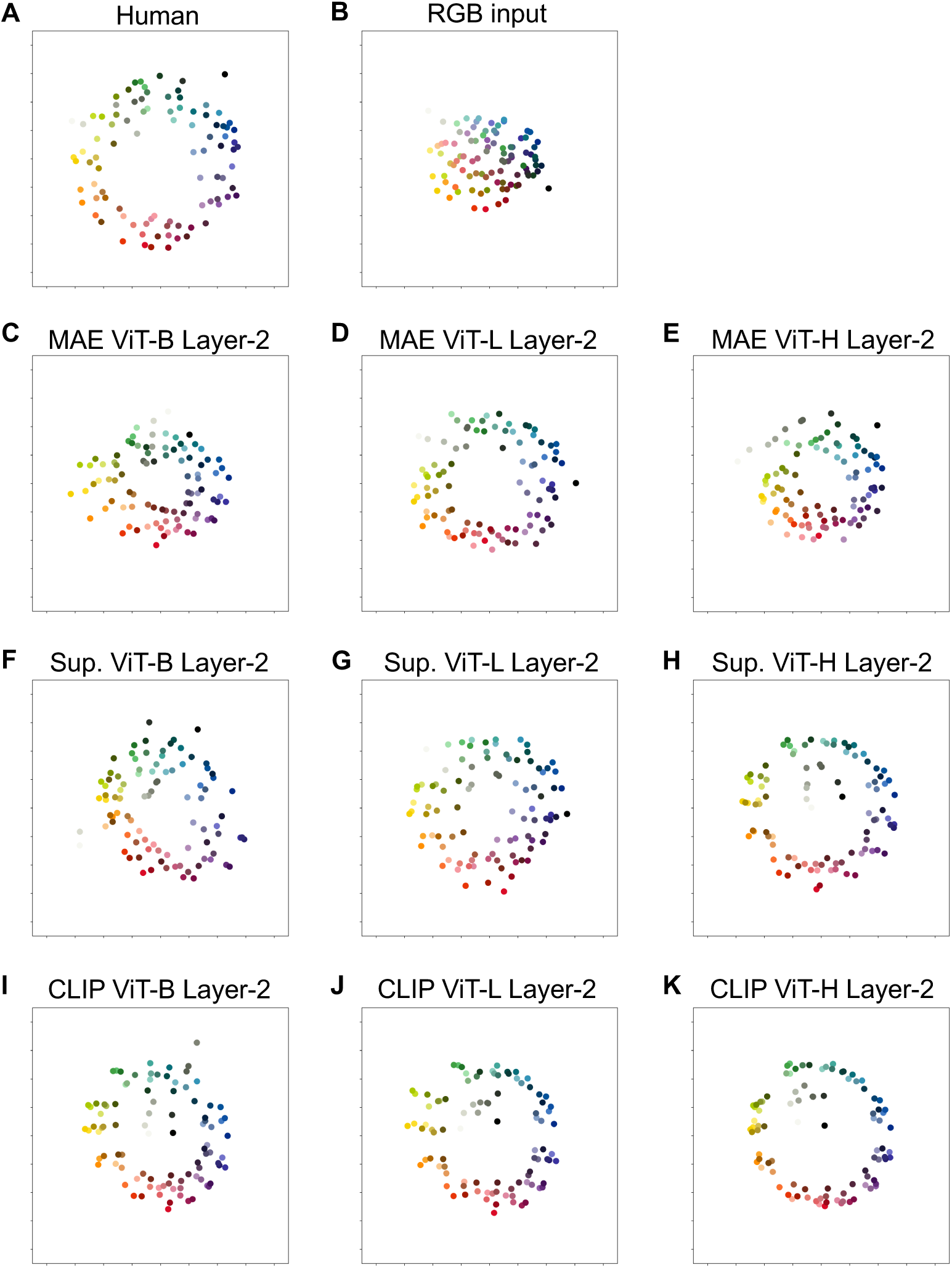
Embeddings of 93 colors from humans, RGB input, and the second-layer embeddings of the DNN models. (A)-(K) Visualization of the embeddings of 93 colors by applying 2-dimensional MDS on the dissimilarity matrix from (A) the color-neurotypical participants group, (B) the RGB model, (C)-(E) the MAE models, (F)-(H) the SL models, and (I)-(K) the CLIP models.

**Fig S10.**
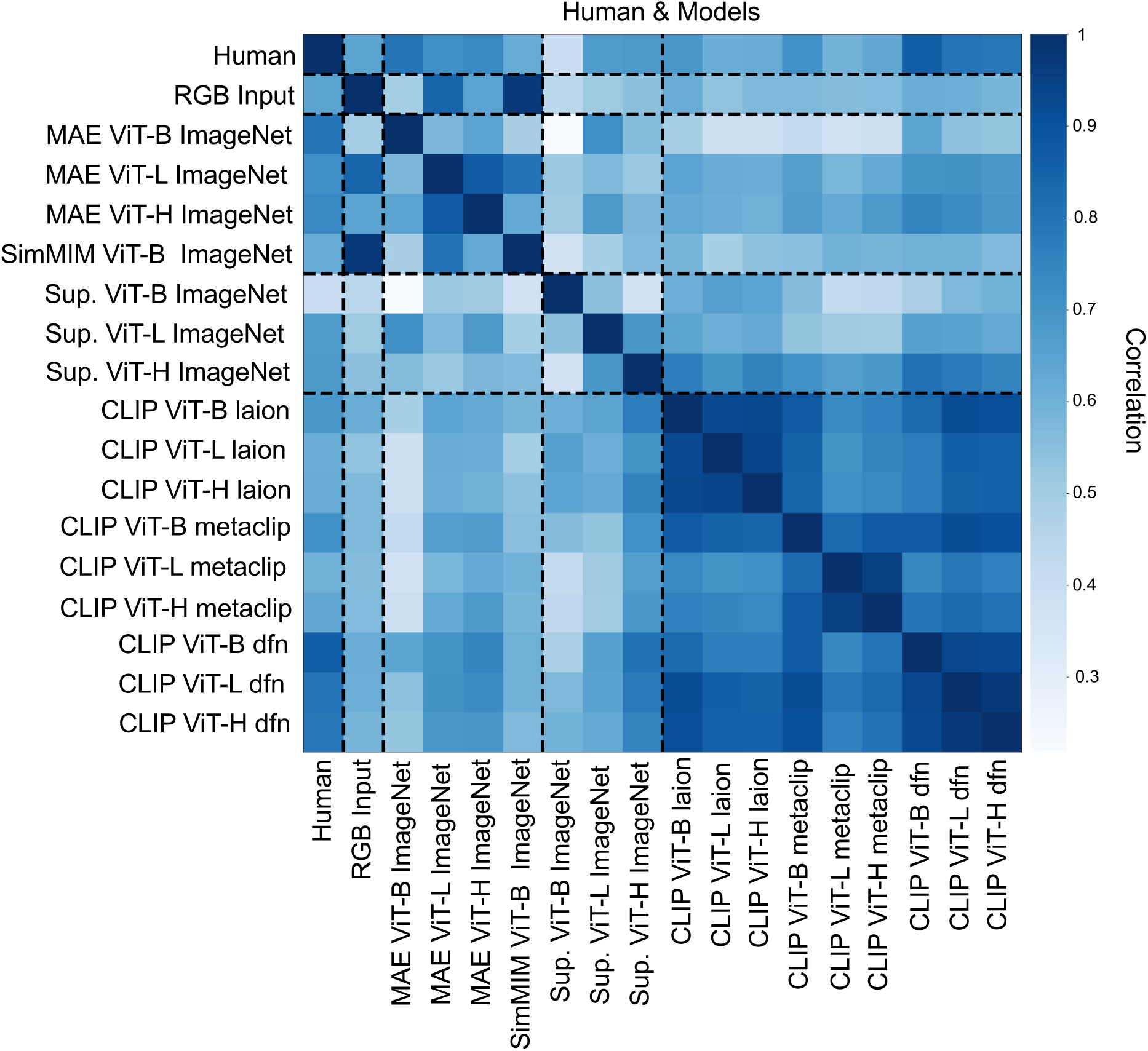
Supervised comparison of color similarity structure for 93 colors between each pair of humans, RGB input, and all of the output embeddings of Transformer-based DNN models. Supervised comparison based on Pearson correlation between each pair of humans, RGB input, and DNN models.

**Fig S11.**
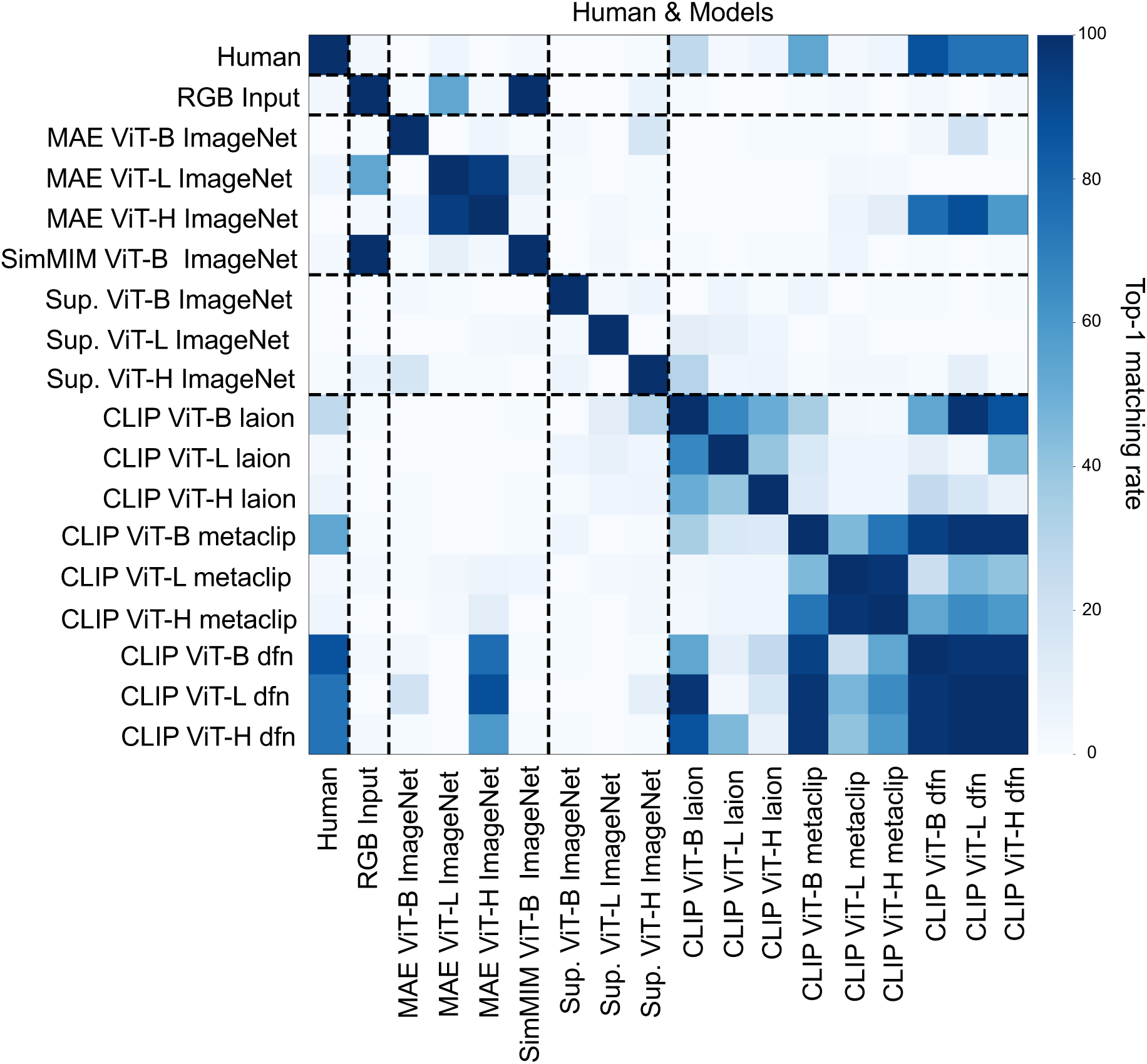
Unsupervised comparison of color similarity structure for 93 colors between each pair of humans, RGB input, and all of the output embeddings of Transformer-based DNN models, with 5 × 10^6^ initializations of GWOT. Unsupervised comparison based on GWOT between each pair of humans, RGB input, and DNN models.

**Fig S12.**
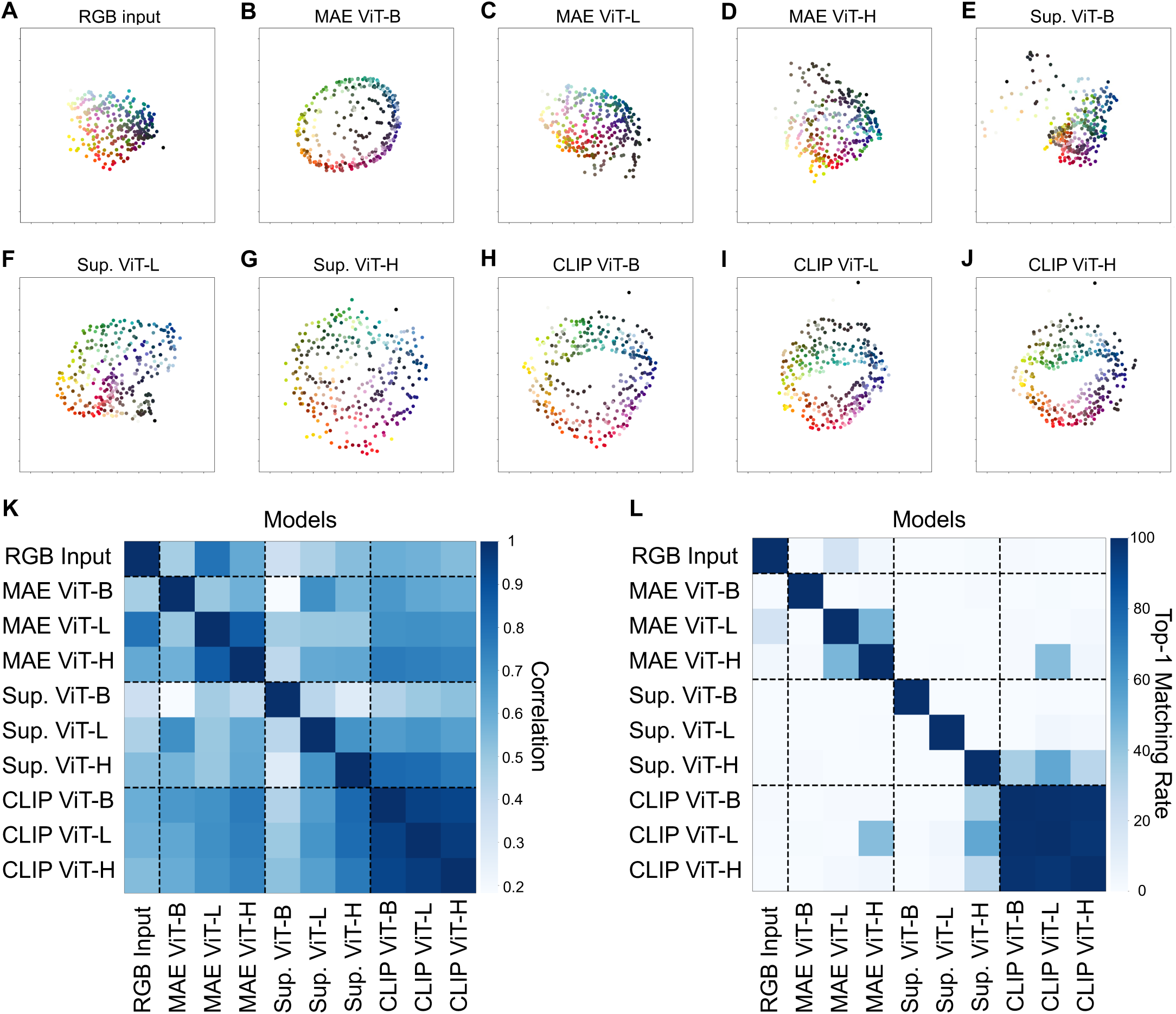
Qualitative and quantitative analysis on 297-color similarity structure of RGB input and the output embeddings of the DNN models. (A)-(J) Visualization of the embeddings of 297 colors by applying 2-dimensional MDS on the dissimilarity matrix from (A) the RGB input, (B)-(D) the MAE models, (E)-(G) the SL models, and (H)-(J) the CLIP models. (K) Supervised comparison based on Pearson correlation between each pair of RGB input and DNN models. (L) Unsupervised comparison based on GWOT between each pair of RGB input and DNN models.

**Fig S13.**
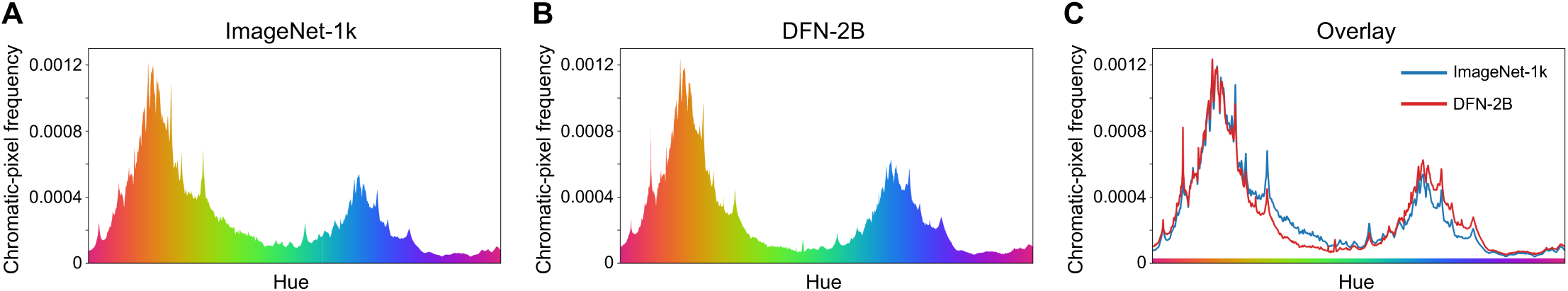
Hue distributions of the datasets used to train the DNN models. (A) Hue distribution of ImageNet-1k (1,281,167 images), used to train the MAE and SL models. (B) Hue distribution of a subset of DFN-2B (1,016,458 images), used to train the CLIP models in the main results. (C) Overlay of the distributions in (A) and (B). The Pearson correlation between the two distributions was *ρ* = 0.97.

**Video S1. Embeddings of 93 colors from humans and RGB input, visualized in three dimensions.** Visualization of the embeddings of 93 colors by applying 3-dimensional MDS on the dissimilarity matrix from the color-neurotypical participants group and the RGB model.

**Video S2. Embeddings of 93 colors from the second-layer embeddings of the DNN models, visualized in three dimensions.** Visualization of the embeddings of 93 colors by applying 3-dimensional MDS on the dissimilarity matrix from the second-layer embeddings of the MAE, SL, and CLIP models.

**Video S3. Embeddings of 93 colors from the output embeddings of the DNN models, visualized in three dimensions.** Visualization of the embeddings of 93 colors by applying 3-dimensional MDS on the dissimilarity matrix from the output embeddings of the MAE, SL, and CLIP models.

**Video S4. Embeddings of 4096 colors from RGB input, visualized in three dimensions.** Visualization of the embeddings of 4096 colors by applying 3-dimensional MDS on the dissimilarity matrix from the RGB model.

**Video S5. Embeddings of 4096 colors from the second-layer embeddings of the DNN models, visualized in three dimensions.** Visualization of the embeddings of 4096 colors by applying 3-dimensional MDS on the dissimilarity matrix from the second-layer embeddings of the MAE, SL, and CLIP models.

**Video S6. Embeddings of 4096 colors from the output embeddings of the DNN models, visualized in three dimensions.** Visualization of the embeddings of 4096 colors by applying 3-dimensional MDS on the dissimilarity matrix from the output embeddings of the MAE, SL, and CLIP models.

## Acknowledgments

We thank Aozora Matsuda for the anlaysis on preliminary stage of this study. Masafumi Oizumi was supported by JST Moonshot R&D Grant Number JPMJMS2012 and Japan Society for the Promotion of Science, Grant-in-Aid for Transformative Research Areas Grant Number 23H04834.

## Author contributions

**Conceptualization:** Nipun Ravindu Wickramanayaka, Masafumi Oizumi

**Data Curation:** Nipun Ravindu Wickramanayaka

**Formal analysis:** Nipun Ravindu Wickramanayaka

**Funding acquisition:** Masafumi Oizumi

**Investigation:** Nipun Ravindu Wickramanayaka

**Methodology:** Nipun Ravindu Wickramanayaka, Masafumi Oizumi

**Project administration:** Masafumi Oizumi

**Software:** Nipun Ravindu Wickramanayaka

**Supervision:** Masafumi Oizumi

**Validation:** Nipun Ravindu Wickramanayaka, Masafumi Oizumi

**Visualization:** Nipun Ravindu Wickramanayaka

**Writing - original draft:** Nipun Ravindu Wickramanayaka, Masafumi Oizumi

**Writing - review & editing:** Nipun Ravindu Wickramanayaka, Masafumi Oizumi

